# Folding heterogeneity in the essential human telomerase RNA three-way junction

**DOI:** 10.1101/2019.12.14.876565

**Authors:** Christina Palka, Nicholas M. Forino, Jendrik Hentschel, Rhiju Das, Michael D. Stone

## Abstract

Telomeres safeguard the genome by suppressing illicit DNA damage responses at chromosome termini. In order to compensate for incomplete DNA replication at telomeres, most continually dividing cells, including many cancers, express the telomerase ribonucleoprotein (RNP) complex. Telomerase maintains telomere length by catalyzing de novo synthesis of short DNA repeats using an internal telomerase RNA (TR) template. TRs from diverse species harbor structurally conserved domains that contribute to RNP biogenesis and function. In vertebrate TRs, the conserved regions 4 and 5 (CR4/5) fold into a three-way junction (3WJ) that binds directly to the telomerase catalytic protein subunit and is required for telomerase function. We have analyzed the structural properties of the human TR (hTR) CR4/5 domain using a combination of in vitro chemical mapping, endogenous RNP assembly assays, and single-molecule structural analysis. Our data suggest that a functionally essential stem loop within CR4/5 is not stably folded in the absence of the telomerase reverse transcriptase protein subunit in vitro. Rather, the hTR CR4/5 domain adopts a heterogeneous ensemble of conformations. RNA structural engineering intended to bias the folding landscape of the hTR CR4/5 demonstrates that a stably folded 3WJ motif is necessary but not sufficient to promote assembly of a functional RNP complex. Finally, single-molecule measurements on the hTR CR4/5 domain show that RNP assembly selects for a conformation that is not the major population in the heterogeneous free RNA ensemble, suggesting that non-canonical hTR folds may be required during telomerase biogenesis.

## Introduction

The ends of linear chromosomes in eukaryotic cells terminate with repetitive DNA sequences that bind to specialized proteins to form telomeres (Blackburn and Gall 1978; Erdel et al. 2017). Telomeres protect coding DNA from degradation and distinguish chromosomal termini from double-stranded breaks in order to evade unwanted recognition by DNA damage response machineries (Muller 1938; McClintock 1939; de Lange 2018). With each round of cell division, the inability of the conventional replication machinery to completely copy the lagging strand template results in gradual telomere attrition. Ultimately the presence of a critically short telomere drives cells into permanent cell growth arrest or apoptosis (Hayflick 1965; Harley et al. 1990). However, cells that must retain high proliferative capacity maintain telomere length through the action of the telomerase reverse transcriptase (Greider and Blackburn 1985; Greider and Blackburn 1989; Kolquist et al. 1998; Wright et al. 2001; Roth et al. 2003). Given the importance of maintaining telomere length in dividing cells, germ-line mutations in telomerase genes result in severe developmental defects (Yamaguchi et al. 2003; Vulliamy and Dokal 2008; Savage 2014). In addition, telomerase contributes to the unchecked cell growth that is a hallmark of human cancers (Kim et al. 1994; Blasco 2005). Therefore, efforts to better understand telomerase structure, function, and regulation have direct biomedical significance.

Telomerase is a multi-subunit ribonucleoprotein (RNP) complex that includes the catalytic telomerase reverse transcriptase (TERT) protein, telomerase RNA (TR), and several additional species-specific holoenzyme proteins that are necessary for proper RNP biogenesis (Egan and Collins 2012a; Chan et al. 2017). The TERT domain architecture is well-conserved across species and consists of the telomerase essential N-terminal (TEN) domain, the telomerase RNA binding domain (TRBD), the reverse transcriptase (RT) domain, and the C-terminal extension (CTE) (Fig. 1A). In contrast, comparison of TRs across species ranging from yeasts to human reveals an exceedingly high degree of variation in both RNA length and sequence (Romero and Blackburn 1991; Chen et al. 2000; Chen and Greider 2004). Interestingly, in spite of this apparent evolutionary divergence, several conserved TR structural elements exist that are essential for enzyme assembly and function. These include the highly conserved template/pseudoknot (t/PK) domain and a stem-terminal element (STE) (Fig. 1B). In vertebrate TRs, the STE is thought to fold into an RNA three-way junction (3WJ) often referred to as the conserved regions 4/5 (CR4/5) domain (Fig. 1C). With regard to TR primary sequence, the CR4/5 domain is spatially separated from the RNA template that must necessarily reside in the TERT enzyme active site; yet, naturally occurring mutations in human telomerase RNA (hTR) CR4/5 can result in human diseases characterized by loss of telomerase function (Yamaguchi et al. 2003; Vulliamy and Dokal 2008; Alder et al. 2018).

**Figure 1:**
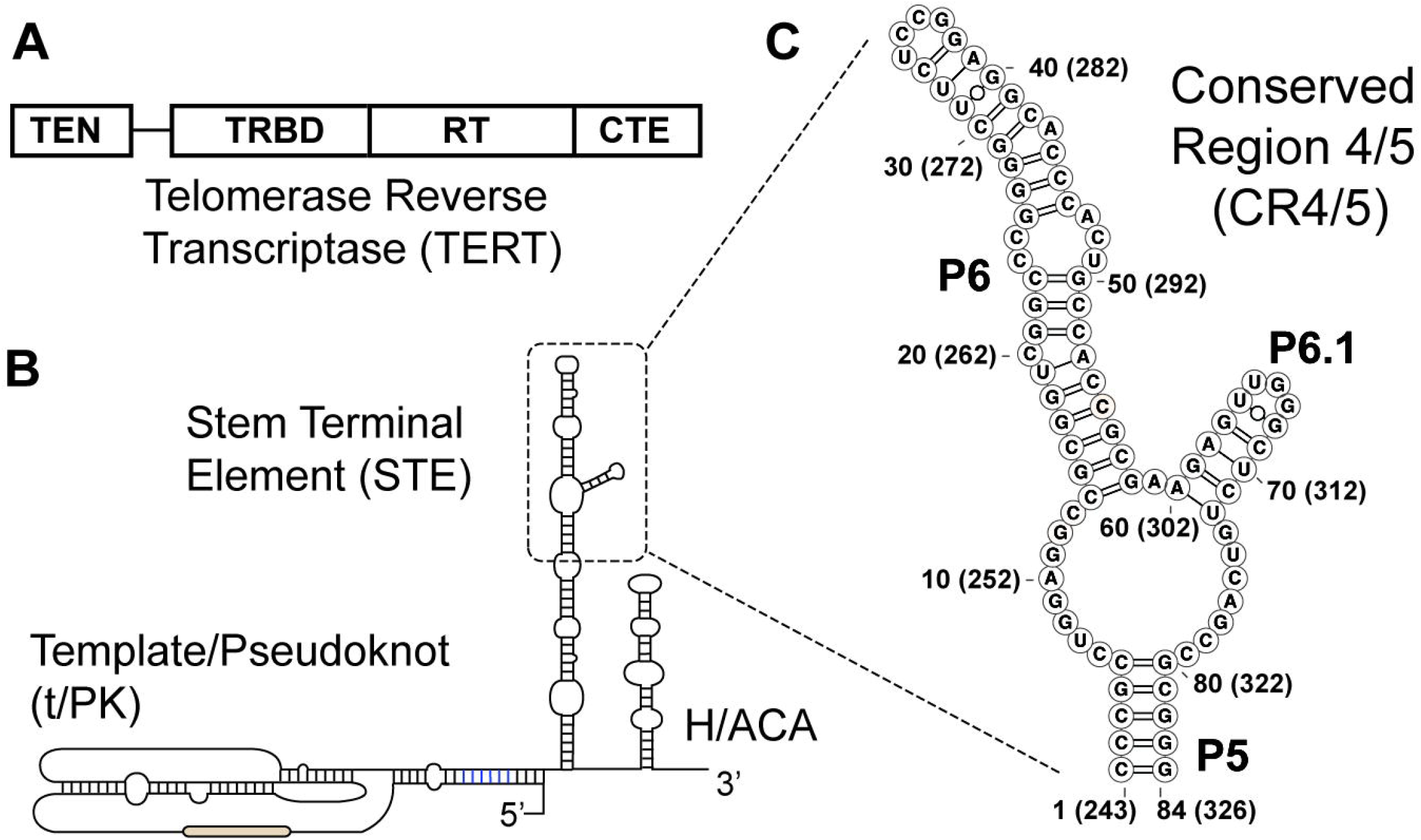
Conserved protein and RNA domains of the telomerase catalytic core. (**A**) The conserved domain architecture of the telomerase reverse transcriptase (TERT) catalytic protein subunit, including: the telomerase essential N-terminal (TEN) domain, the RNA binding domain (TRBD), the reverse transcriptase (RT) domain, and the C-terminal extension (CTE). (**B**) The conserved domain organization of the human telomerase RNA (TR), including the template/pseudoknot (t/PK) domain, the stem terminal element (STE), and the H/ACA box motif. (**C**) Conserved regions 4 and 5 (CR4/5) domain of the human TR (hTR) comprised of stems P5, P6, and P6.1. Nucleotide numbering system used throughout the study is indicated together with the corresponding nucleotide numbering within full-length hTR in parentheses.

In hTR, the CR4/5 domain includes three RNA helices (P5, P6, and P6.1) joined together by an expanded RNA junction sequence (Fig. 1C). Detailed biochemical studies performed on vertebrate TR CR4/5 variants have shown that a stably formed P6.1 helix within the 3WJ is essential for telomerase assembly and function (Mitchell and Collins 2000; Chen et al. 2002; Kim et al. 2014). Chemical and enzymatic RNA structure probing experiments of full length hTR have led to mixed conclusions as to the structural state(s) of the essential P6.1 stem loop within the CR4/5 domain prior to and upon telomerase assembly (Antal et al. 2002; Zemora et al. 2016). More recently, the human telomerase holoenzyme protein TCAB1 was implicated in mediating a conformational change in the CR4/5 3WJ domain (Chen et al. 2018). Protein-RNA crosslinking studies and an atomic-resolution structure of the medaka fish TR 3WJ bound by its cognate TERT-TRBD revealed the molecular details of the TERT-RNA interaction (Bley et al. 2011; Kim et al. 2014). Interestingly, the helical arrangement observed in the medaka protein-RNA complex was substantially altered when compared to the solution structure of the same RNA domain in the absence of protein (Huang et al. 2014). Over the last several years, cryo EM structures of the *Tetrahymena* and human telomerase RNPs were reported (Jiang et al. 2018; Nguyen et al. 2018), providing additional details on the arrangement of protein and RNA domains within the fully assembled telomerase RNP complex. Both structures suggest that an apical stem loop within the STE (P6.1 in hTR) lies at the interface of the TERT-CTE and TERT-TRBD domains, providing clues as to the essential requirement of the P6.1 stem loop in coupling the two TERT domains during telomerase assembly and/or function.

Here, we set out to characterize the in vitro RNA folding properties of the hTR CR4/5 domain using a combination of chemical mapping, paired together with single-molecule Förster Resonance Energy Transfer (smFRET) experiments. Chemical probing experiments using a variety of RNA modification reagents revealed a substantial degree of reactivity within the region of hTR CR4/5 expected to form the essential P6.1 stem loop structure. Use of chemical reactivity data to guide computational modeling of RNA structure predicts several possible alternative conformations that may be sampled within the hTR CR4/5 structure ensemble. To further validate these structure predictions, we systematically perturbed each nucleotide within the hTR CR4/5 domain, and queried the effects of each mutation on the chemical reactivity profile (Kladwang et al. 2011a; Tian et al. 2014). The results of these Mutate-and-Map (M^2^) experiments reinforce the conclusion that the P6.1 stem loop is not well ordered in vitro. Next, we engineered hTR CR4/5 variants designed to bias the folding energy landscape to favor P6.1 formation. After validating the efficacy of the RNA designs by chemical probing in vitro, selected full-length hTR constructs were transfected into human cells to be endogenously assembled and purified. We find that certain hTR sequence variants intended to stabilize the canonical P6.1 fold displayed marked defects in assembly of functional RNP complexes, demonstrating that a stably folded hTR 3WJ is necessary but not sufficient for telomerase function. Using smFRET to probe the conformational properties of the hTR CR4/5 domain also revealed heterogeneous RNA folding, characterized by at least three distinct FRET states. Interestingly, smFRET measurements made on the same hTR CR4/5 fragment assembled into an active RNP complex show that the conformation of the RNA is substantially altered upon protein binding. Collectively, our results are consistent with a working model wherein alternative hTR folds may serve as RNP assembly intermediates during telomerase biogenesis, as has been shown for other essential cellular RNPs such as the ribosome and spliceosome (Abelson et al. 2010; Mohan and Noller 2017). Given the central importance of CR4/5 folding in promoting telomerase assembly and function, the hTR folding properties characterized in the present work provide a new framework for developing small molecules that target hTR assembly intermediates with the goal of inhibiting telomerase in cancer cells.

## RESULTS

### Chemical probing of the telomerase RNA three-way junction

The 3WJ motif is well conserved across many telomerase RNA systems, ranging from yeasts to vertebrates. Many of the RNA structural models that are used to generate hypotheses relating to telomerase function are derived from sequence covariation analysis (Chen and Greider 2004) and/or the use of biochemical mutagenesis (Mitchell and Collins 2000; Chen et al. 2002). One challenge of methods such as sequence covariation analysis is that the resultant models may not accurately capture the structural properties of all RNA folding intermediates before it interacts with physiological binding partners. Indeed, studies of telomerase biogenesis indicate that hTR accumulates in sub-nuclear compartments prior to assembly with the TERT protein subunit (Etheridge et al. 2002; Zhu et al. 2004), raising the distinct possibility that hTR may exist in various structural states prior to telomerase assembly. In order to better understand the structural properties of TRs prior to and during RNP biogenesis, we set out to analyze the secondary structural properties of telomerase 3WJs from two vertebrate systems: medaka fish (*Oryzias latipes*) and human. The medaka TR 3WJ serves as an important benchmark in our TR structural analyses because its atomic structure is well characterized in the absence and presence of the TERT-TRBD (Huang et al. 2014; Kim et al. 2014).

For each TR system, we used an isolated CR4/5 RNA fragment to facilitate in vitro structure probing. Notably, the isolated hTR CR4/5 domain used in our studies is sufficient to support telomerase function when reconstituted with the hTR t/PK domain and TERT protein (Fig. S1) (Tesmer et al. 1999). Several sequence elements were added to the TR segment to assist in quantitative data analysis of chemical probing experiments (Fig. 2A). First, a primer-binding site was appended to the RNA 3’-end for use in the reverse transcriptase reactions required to read out sites of RNA modification. Second, a short RNA hairpin structure flanked by unstructured ‘buffer’ regions was added to serve as an internal normalization control when calculating chemical reactivities (see Methods for details) (Kladwang et al. 2014). De novo structure predictions using only the RNA sequences as calculated on the RNAstructure web server (Reuter and Mathews 2010) yielded lowest free energy conformations with the expected stems that collectively form the 3WJ fold (Fig. 2B). In the case of the hTR CR4/5 domain, RNAstructure predicted an additional cross-junction clamping helix not typically included in canonical representations of this region of hTR. Furthermore, multiple structures with nearly isoenergetic stability were also predicted, including conformations lacking the essential P6.1 stem loop (Fig. S2), highlighting the need for experimental data to validate specific RNA models.

**Figure 2:**
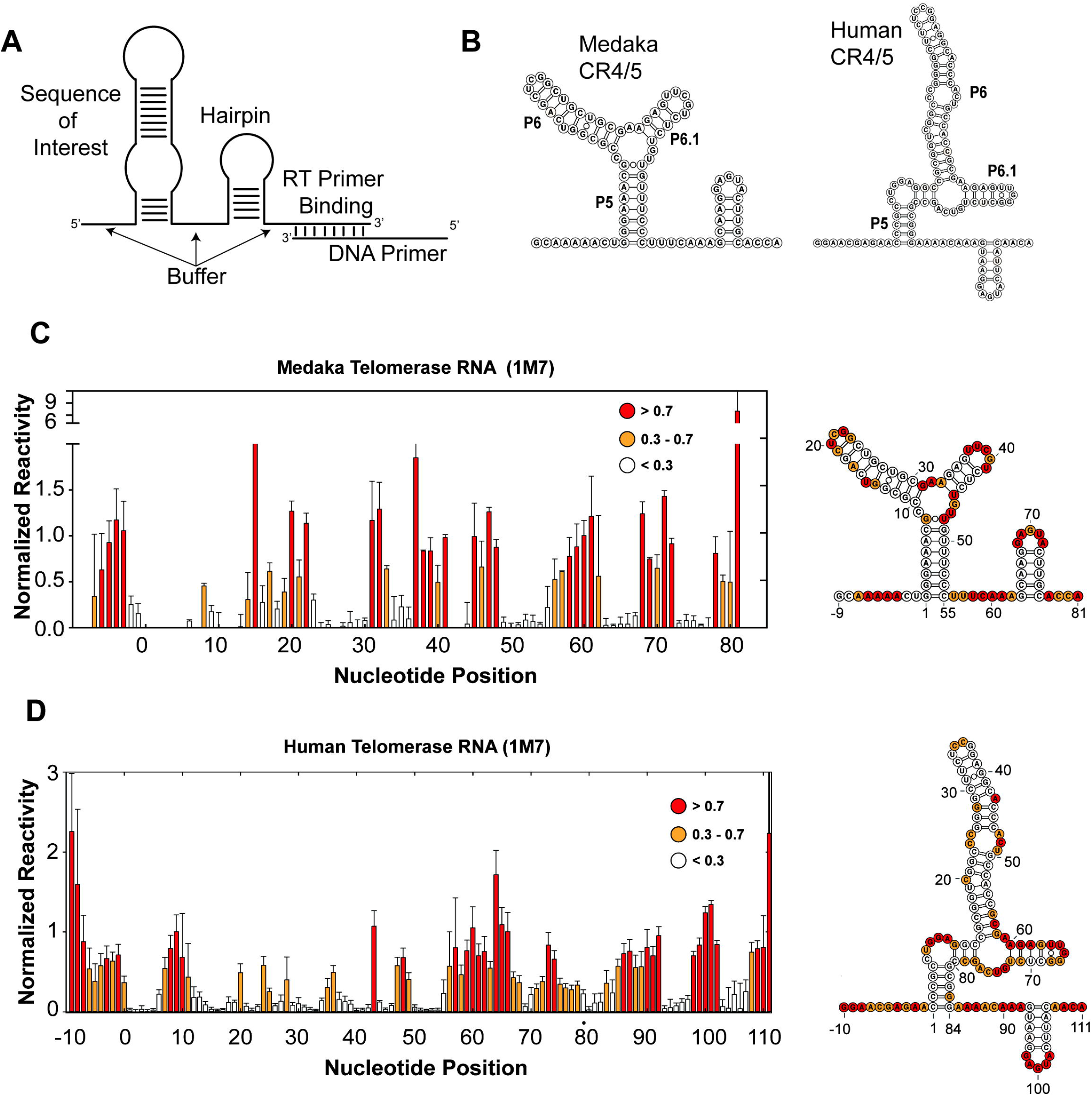
Chemical mapping of medaka and human CR4/5 domains. (**A**) Cartoon schematic of general RNA construct design, including the RNA sequence of interest flanked by unstructured RNA buffer sequences, a normalization RNA hairpin, and a reverse transcriptase priming site. (**B**) Lowest energy predicted secondary structure of medaka (left) and human (right) CR4/5 domain using RNAstructure. (**C**) (left) Chemical mapping of the medaka CR4/5 domain by SHAPE (1M7 probing). Plotted normalized reactivity values are color-coded (red > 0.7, yellow 0.3 - 0.7, and white < 0.3). Each bar plotted represents experiments conducted in triplicate with the respective standard deviation as error bar. (right) Color-coded schematic of the reactivity data is shown on the RNAstructure predicted secondary structure. (**D**) Chemical mapping of the human CR4/5 domain by SHAPE. Color coding in bar plot and structure schematic is as described in panel C.

To experimentally evaluate each of these CR4/5 structure predictions we performed selective hydroxyl acylation analyzed by primer extension (SHAPE) experiments using 1-methyl-7-nitoisatoic anhydride (1M7), a fast acting chemical modifier (Mortimer and Weeks 2007). The 1M7 SHAPE reagent acylates the 2’-hydroxyl of flexible ribose moieties along the RNA backbone, thereby providing a sequence-independent proxy for unstructured regions of an RNA. For each of these ‘chemical mapping’ experiments, sites of chemical modification were read out as premature termination sites during RT catalyzed primer extension. The individual 1M7 reactivity at every position along the RNA was calculated and then normalized to the internal hairpin control signal (see Methods for details). In addition, experiments were also performed using the base-specific reagents dimethyl sulfate (DMS) or 1-cyclohexyl-(2-morpholinoethyl)carbodiimide metho-*p*-toluene sulfonate (CMCT), which primarily react with adenine/cytosine or guanine/uracil bases, respectively (Fig. S3). In the case of the medaka TR CR4/5 domain, reactivity profiles obtained by all three chemical probing methods (DMS, CMCT, and 1M7) yielded data that support the canonical base pairing arrangement expected for this 3WJ fold, and are highly consistent with the reported solution structure of this same RNA fragment (Kim et al. 2014) (Fig. 2C and Fig. S3 and S4). In contrast, strong 1M7 reactivity was observed in the region of the hTR CR4/5 domain expected to fold into the P6.1 stem (Fig. 2D). This result is unexpected given the established importance of the P6.1 stem loop structure in promoting telomerase RNP assembly and function (Mitchell and Collins 2000; Chen et al. 2002; Kim et al. 2014). Taken together, these data suggest that using sequence information alone, the RNAstructure folding algorithm can effectively capture the expected base pairing configuration of the medaka TR 3WJ, but fails to do so in the more expanded junction/P6.1 region of the hTR CR4/5 domain.

### SHAPE-guided modeling of telomerase RNA CR4/5 domain structure

RNAstructure calculates the lowest free energy structures using thermodynamic parameters that are dynamically sampled against databases of structures with well-characterized stabilities (Reuter and Mathews 2010). Experimentally derived chemical probing data significantly improves the predictive power of the RNAstructure folding algorithm (Mathews et al. 2004). For example, SHAPE reactivities are typically used to generate a pseudo-energy change term (ΔG_SHAPE_) for each individual base pair of a predicted structure, which can then be added to the RNAstructure prediction algorithm as a nearest neighbor free energy term (Deigan et al. 2009). Using this approach, we generated SHAPE-guided models of the medaka TR and the hTR CR4/5 domains with the Biers component of the HiTRACE software package that implements the use of the RNAstructure prediction algorithm (Kladwang et al. 2011a; Tian et al. 2014). In addition to predicting a lowest energy conformation for a particular RNA sequence utilizing SHAPE reactivity data, Biers also includes a nonparametric bootstrapping function to estimate confidence levels in the prediction of each helical element within a particular low energy RNA conformation (Kladwang et al. 2011b). Specifically, the bootstrapping function within the Biers software package uses random resampling with replacement of experimental SHAPE data to calculate a collection of data-guided RNAstructure outputs (Fig. S5). This collection of bootstrapping-derived structures is then used to calculate the frequency of each base pair present across all computationally derived replicates. In this way, the resulting bootstrap value for any given helix provides a metric to help evaluate the degree of confidence for each helix given the data that were used to guide the structure calculation. It is important to note that bootstrap values are a statistical tool to analyze computational prediction methods, and should not be interpreted as an indicator of the equilibrium conformation(s) present for a particular RNA of interest.

As expected, the addition of the ΔG_SHAPE_ constraints to predictions of the medaka TR CR4/5 yields the canonical 3WJ fold with each of the expected helices being called with high bootstrap values (Fig. 3A). We note that structure prediction was performed on the complete RNA construct, including the normalization hairpin which was also predicted with high confidence. This result indicated that addition of experimentally derived data does not cause the RNAstructure algorithm to deviate in its prediction of the lowest energy conformation for the medaka TR CR4/5. In the case of the hTR CR4/5, the inclusion of ΔG_SHAPE_ constraints in structure calculation recaptures a lowest energy conformation in which the P5, P6, and normalization hairpin are called with high confidence. In contrast, the bootstrap value calculated for the P6.1 stem is significantly decreased, consistent with the high levels of SHAPE reactivity in this region (Fig. 3B). Manual inspection of alternative predicted RNA conformations with nearly equal calculated energies yield a substantially altered junction region defined by extensive base pairing that is not characteristic of a 3WJ fold (Fig. 3B). These data-driven structure predictions indicate the hTR CR4/5 domain is structurally heterogeneous within the expanded junction region and the functionally essential P6.1 stem loop is not stably folded in vitro.

**Figure 3:**
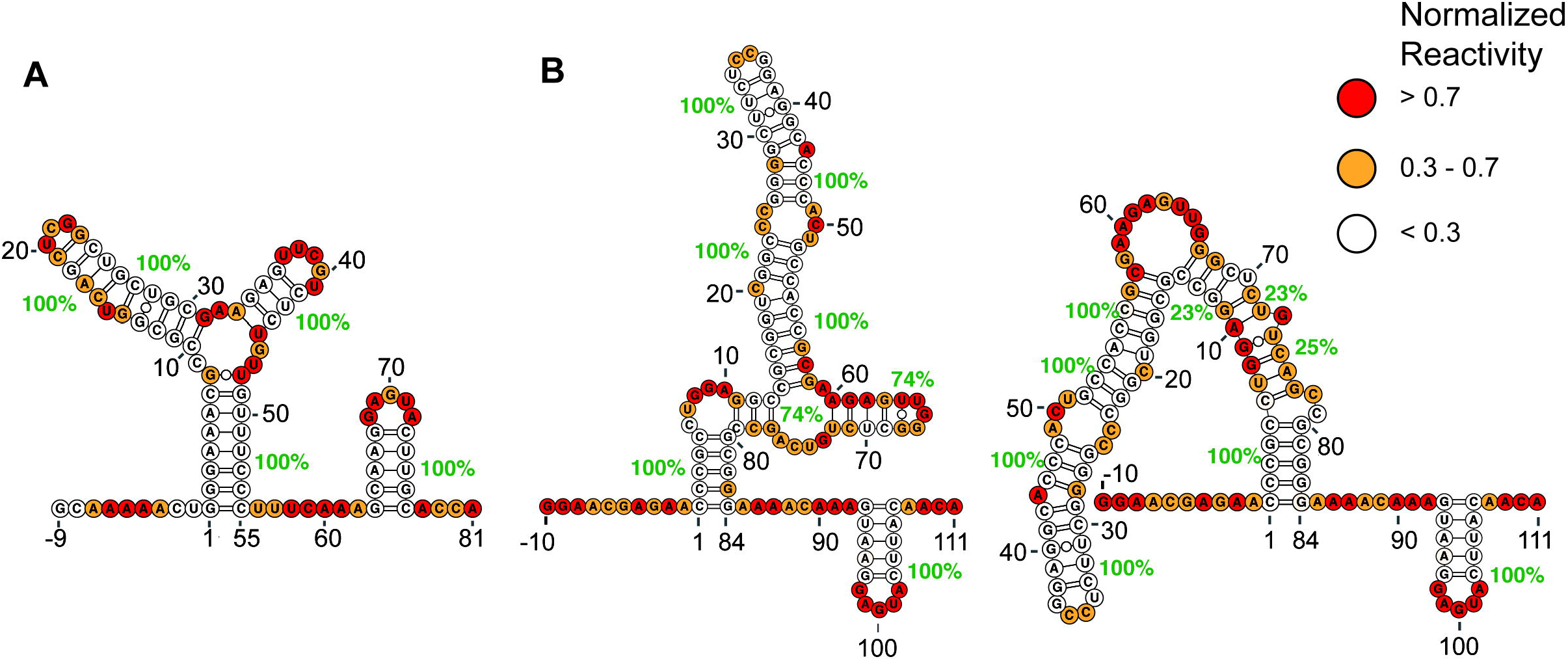
Data guided RNA secondary structure prediction of medaka and human CR4/5 domain. SHAPE (1M7) reactivity data was as used as weights to guide RNA structure prediction for medaka (**A**) and human (**B**) CR4/5 domains. Green percentage values indicate bootstrap support levels for each helical element in the predicted structures. Bootstrap analysis for each data set was used to estimate variation in the predicted structures across each helix. Normalized reactivity values are indicated on each of the data-guided RNA structures (red > 0.7, yellow 0.3 - 0.7, and white < 0.3).

### Multi-dimensional chemical mapping supports hTR CR4/5 structural heterogeneity

To further probe the structure of the hTR CR4/5 domain, we performed multi-dimensional chemical mapping (MCM) (Kladwang et al. 2011a). This systematic mutagenesis approach permits rapid chemical probing analysis of a panel of RNA mutant constructs designed to explicitly test for the presence of Watson-Crick base pairing in a proposed RNA secondary structural model (Kladwang et al. 2011a; Tian et al. 2014). If a mutation is made to a base that is engaged in a base pair, then one expects the release of the interacting partner that consequently becomes accessible to the SHAPE probe. In order to probe for such specific release events, we generated a set of 84 mutants across the entire hTR CR4/5 construct. The chemical reactivity profiles of all RNA variants were stacked vertically to generate a reactivity tapestry (Fig. 4A). Signals on the diagonal of the reactivity tapestry represent release events at the engineered site of mutation (Fig. 4A, red dotted line). Signals that deviate from the wild type reactivity profile indicate changes in reactivity that result from each individual mutation. Many of the single mutant reactivity profiles revealed complex structural rearrangements beyond the simple base pair release event principle. However, visual inspection of the data reveals multiple features in the reactivity tapestry that support specific base pairs present within the hTR CR4/5 (Fig. 4A, red circles, and Fig. 4B). For example, the G27C and G28C mutations each resulted in increased reactivity at positions C45 and C44, respectively, providing support for these base pairs being present within the P6b stem (Fig. 4B). Similarly, the C44G and C45G mutations resulted in release events in G28 and G27, respectively, providing independent support for these same base pairs in the P6b stem. Increased reactivity was also observed for certain mutations within the P6a stem; for example, the C51G, C54G, and G56C mutations each caused increased signal at positions G22, G18, and C16, respectively. Lastly, the G82C mutation located within the P5 stem resulted in increased reactivity at position C3, providing support for this specific base pairing interaction. Notably, the high baseline reactivity observed in the hTR CR4/5 junction and P6.1 stem loop region precludes unambiguous visual analysis of the MCM data. However, we found that mutations introduced at the base of the P6a stem (A53U, C55G, and C57G) had the unexpected effect of causing substantial structural rearrangements in the CR4/5 domain, evidenced by reduced reactivity in the junction region and increased reactivity within the P6 stem (Fig. 4A, blue arrows). Other notable global folding changes were observed for single G→C substitutions located within the P6.1 stem, such as G61C and G63C, which both induce the CR4/5 domain to fold into an extended two-helix junction (Fig. 4A, purple arrows, and Fig. S6).

**Figure 4:**
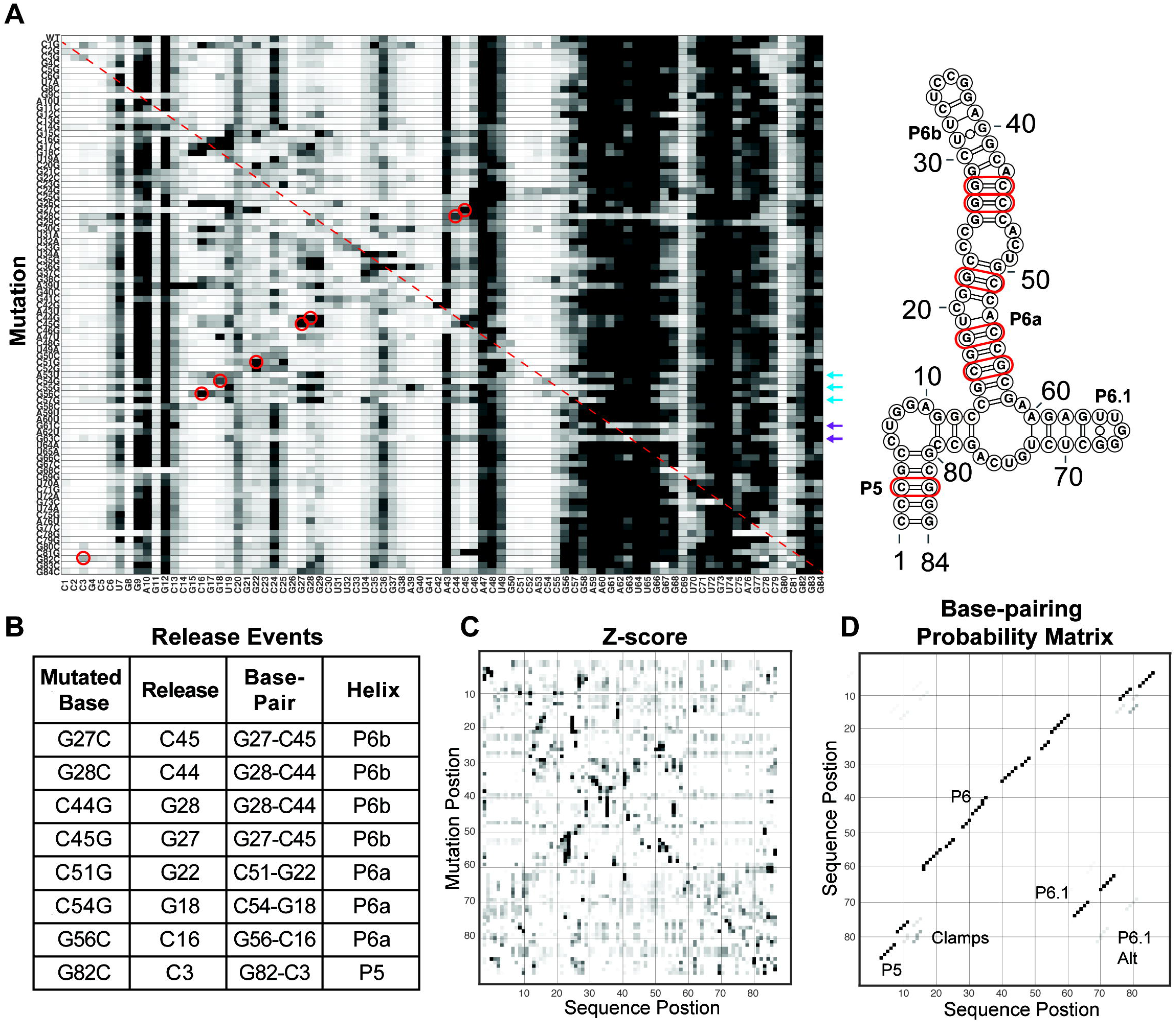
Mutate-and-map profiling of the human CR4/5 domain indicates presence of structural heterogeneity within the RNA junction region. (**A**) Systematic mutations were introduced at each base within the hTR CR4/5 domain as indicated ((A → U, U → A, G → C and C → U). The structure of each mutant was interrogated by SHAPE (1M7) and the resultant reactivity profiles were stacked to create a reactivity tapestry that permits visual comparison of the chemical reactivity at each nucleotide across all mutants. Red dashed line corresponds to position of expected signal of enhanced reactivity at the site of the base substitution. Specific sites of enhanced reactivity (‘release events’) are circled in red. Positions of validated base pairing interactions are highlighted in red in the secondary structure model shown to the right. Mutation positions with the P6 stem (blue arrows) and P6.1 stem (purple arrows) that induce large-scale changes in the reactivity patterns are indicated. (**B**) Summary of specific mutations and sites of correlated enhancements of chemical reactivity together with the positions of the CR4/5 base pairs that these data support. (**C**) A Z-score plot indicating patterns of mutation-induced changes in chemical reactivity at each nucleotide position across all RNA mutants. The Z-scores are used as weights to guide structure prediction in the RNAstructure software. (**D**) Bootstrap support values are plotted in a base pair probability matrix represented in grey scale. High confidence stems give rise to dark and symmetric signals. Each of the RNA structure elements are annotated including non-canonical cross junction clamps and an alternate P6.1 stem.

To achieve a quantitative analysis across the entire reactivity tapestry we generated a Z-score plot, where individual Z-scores report on the statistical significance of deviations in the reactivity level for a given nucleotide compared across all RNA constructs (Fig. 4C). Z-score values are then used as a pseudo-energy term to guide structure prediction by RNAstructure within the Biers component of the HiTRACE software package (Tian et al. 2014). As with the SHAPE reactivity-guided RNAstructure calculations, the Z-score data can be used to perform bootstrapping analysis as a measure of confidence in each predicted helical segment and to generate a base pair probability matrix (Fig. 4D). The results of the Z-score analysis are consistent with the presence of structures other than the canonical P5, P6 and P6.1 stems in the CR4/5 structure ensemble. For example, in multiple Z-score driven structures an alternative P6.1 stem (P6.1 alt) was predicted in addition to several mutually exclusive cross-junction clamping helices (Fig. 4D). Taken together, the results of the MCM experiments provide additional experimental evidence for base pairing interactions in the P6a, P6b, and P5 stems, and support the notion that the junction region and P6.1 stem loop may adopt non-canonical base pairing configurations.

### Increasing the stability of the P6.1 stem loop is not sufficient to support functional telomerase RNP assembly

In order to test the hypothesis that structural heterogeneity within the hTR CR4/5 contributes to telomerase assembly and function, we employed RNA structural engineering to bias the folding properties the 3WJ motif. Both naturally occurring or engineered RNA mutations that disrupt or destabilize the essential P6.1 stem loop result in significant telomerase loss-of-function phenotypes (Mitchell and Collins 2000; Chen et al. 2002; Kim et al. 2014). Here, we focused on RNA mutants intended to improve the folding stability of the P6.1 stem loop. First, we replaced the native hTR 3WJ bulge sequence with poly-A stretches (Mut-1), leaving the P6.1 sequence unaltered. Second, we designed a CR4/5 domain with three mutations (A302C, U306C, U314G) resulting in two additional G-C base pairs within the P6.1 stem, but preserving the native sequence of the expanded junction region and the P6.1 loop (Mut-2). To validate the designs of Mut-1 and Mut-2, we measured the chemical reactivity of each of the variants. In the case of both mutants significantly less reactivity was observed in the region around the P6.1 stem (nt 58-72) when compared to the wild type RNA (Fig. 5A). Apparent SHAPE protection at the 5’ ends of the poly-A segments in Mut-1 (Fig. 5B) is explained by a recently discovered anomaly in the reverse transcription reaction (Wellington-Oguri et al. 2018) (R. Das unpublished results). For both Mut-1 and Mut-2 hTR variants, SHAPE-driven modeling of RNA structures supported the conclusion that the engineered sequence changes stabilized the P6.1 stem loop fold (Fig. 5B).

**Figure 5:**
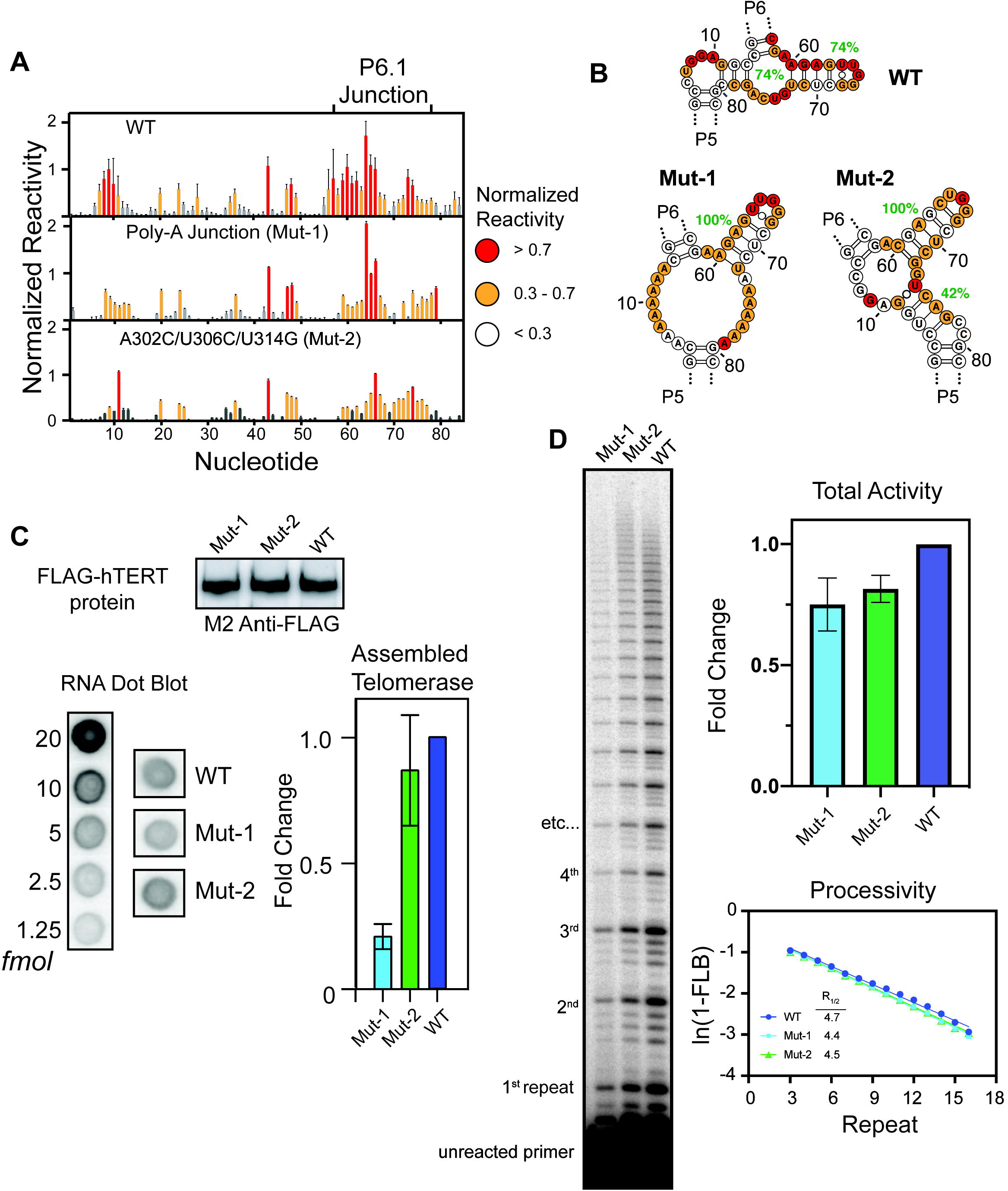
RNA engineering of the CR4/5 domain that stabilizes the P6.1 stem impacts endogeneous telomerase assembly. (A) SHAPE (1M7) reactivity data of wild-type (WT), Poly-A junction (Mut-1), and the A302C/U306C/U314G triple mutant (Mut-2) are plotted as normalized reactivity values and color-coded (red > 0.7, yellow 0.3 - 0.7, and white < 0.3). Bar plot represents experiments conducted in triplicate and error bars are the standard deviation of the three replicates. (**B**) Normalized reactivity values for each of the CR4/5 constructs are mapped onto the data-guided secondary structure prediction of the junction region with bootstrap support values for each helical element shown in green. (**C**) Western blot analysis using an anti-FLAG antibody was performed on each of the immunopurified telomerase complexes (top). RNA dot blot data, together with a set of RNA standards, is shown (lower left). RNA dot blot data were quantified, normalized to the western blot signal, and then plotted as a measure for the amount of CR4/5 domain assembled into telomerase complex and purified from HEK 293T cells (lower right). (**D**) Direct DNA primer extension assay on equal volumes of purified telomerase for each CR4/5 construct (left). The position of the unextended ^32^P-labeled DNA primer as well as each major telomere repeat band is indicated to the left of the gel. Total activity in each lane, normalized for loading differences based upon the total counts per lane, are plotted (top right), and processivity values were extracted by analyzing the decay pattern of the major telomere repeat bands (lower right). Error bars in total activity plot represent standard deviation of experiment performed in triplicate. RAP estimates are taken from the plotted fit of a representative primer extension experiment.

Having validated the Mut-1 and Mut-2 designs in vitro, we next sought to investigate the impact of these same changes within the context of full-length hTR expressed and assembled in HEK 293T cells. FLAG-tagged human TERT protein and each hTR variant were co-expressed after transient transfection and immunoprecipitated for biochemical characterization. Western blots against immunoprecipitated FLAG-TERT demonstrated comparable levels of protein expression and pull-down efficiency across all experiments (Fig. 5C, top panel). Similarly, RNA dot blot analysis was used to analyze the amount of recovered hTR during the FLAG immunoprecipitation procedure. Control experiments indicated that recovery of hTR above background was dependent on the presence of FLAG-TERT (data not shown). The amount of detected hTR was hence used as a measure for efficiency of telomerase assembly. When compared to wild type hTR, the Mut-1 (Poly-A) variant showed a decrease in telomerase assembly (Fig. 5C, bottom). In contrast, the Mut-2 (A302C, U306C, U314G) construct did not exhibit a statistically significant decrease in the efficiency of telomerase assembly (Fig. 5C, bottom). Next, we assayed the endogenously assembled and purified telomerase RNP complexes for catalytic activity using a direct primer extension assay (Fig. 5D). Both Mut-1 and Mut-2 show a modest decrease in the total telomerase activity observed (Fig. 5D). Interestingly, however, estimates of the repeat addition processivity for the variant enzymes indicate that the functional RNP complexes in each case possess similar catalytic properties. Taken together, these results indicate that the presence of a stably folded P6.1 stem loop is not sufficient to promote wild type levels of telomerase assembly in HEK 293T cells. Rather, regions of the expanded junction within the CR4/5 domain appear to be necessary to adopt alternate RNA folds that may be required for telomerase assembly and/or to mediate protein-RNA contacts within the functionally assembled RNP.

### Single-molecule analysis reveals CR4/5 folding heterogeneity and remodeling upon telomerase RNP assembly

Results from our ensemble chemical probing experiments suggest that the human CR4/5 domain exhibits folding heterogeneity, particularly in the junction region that is proximal to the functionally essential P6.1 stem loop. To directly test for folding heterogeneity in the hTR CR4/5 domain we employed a single-molecule Förster Resonance Energy Transfer (smFRET) technique, which measures RNA conformation(s) as the distance-dependent energy transfer between a FRET donor (Cy3) and an acceptor (Cy5) dye incorporated into the RNA. FRET probes were strategically incorporated at positions U32 (Cy5) and U70 (Cy3) to establish a dye pair that reports on the physical proximity of the P6 and P6.1 stem loops (Fig. 6A). Importantly, reconstitution of the fluorescently-labeled CR4/5 domain into human telomerase RNPs using rabbit reticulocyte lysate (RRL) yielded telomerase RNP enzymes that displayed catalytic activity comparable to unlabeled telomerase assessed by primer extension assays (Fig. S1).

**Figure 6:**
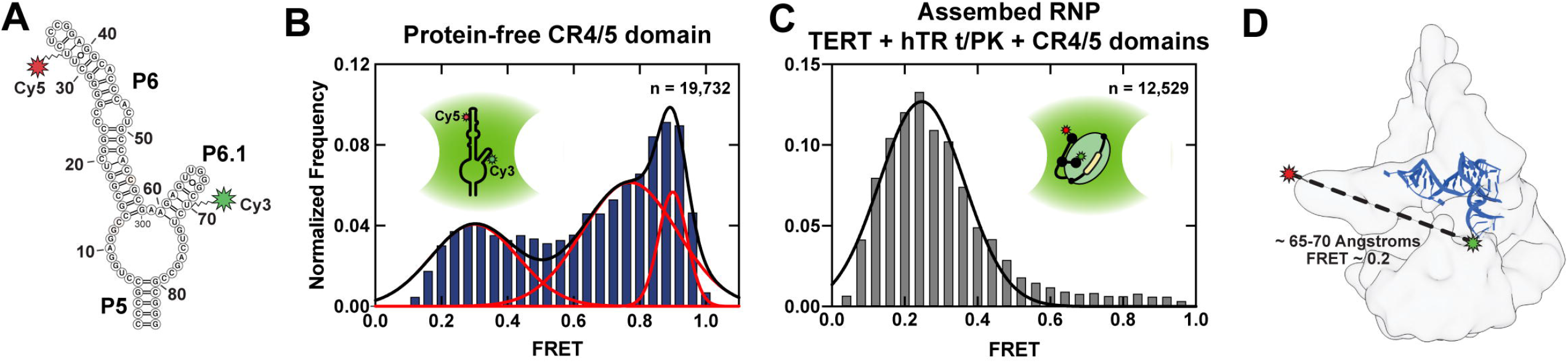
Single-molecule FRET analysis reveals structural heterogeneity in the CR4/5 domain and stabilization of a single population within the assembled telomerase RNP. (**A**) Single-molecule FRET construct design of the CR4/5 domain harboring a donor (Cy3) dye at position U70 and acceptor (Cy5) dye at position U32. (**B**) Data were collected using a single-molecule confocal microscope to measure FRET distributions of the double-labeled CR4/5 domain freely diffusing in solution. Gaussian fitting to the FRET distribution reveals three FRET populations centered at 0.3, 0.77, and 0.9, respectively. The number of independent FRET measurements included in the histogram is indicated and the solution free CR4/5 domain is illustrated schematically. (**C**) Double-labeled CR4/5 was reconstituted with TERT and the template/pseudoknot (t/PK) domain into the telomerase RNP and confocal microscopy was used to measure the FRET distribution. A single FRET population centered at 0.25 was observed. The number of independent FRET measurements included in the histogram is indicated and the telomerase RNP is illustrated schematically. (**D**) Cryo EM density of human telomerase (EMD-7518) (Nguyen et al. 2018) with the medaka CR4/5 crystal structure (blue, derived from the CR4/5-TRBD structure PDB 4O26) (Huang et al. 2014) manually docked. Approximate locations of each FRET dye are indicated and the distance between these positions within the structural model is indicated together with the estimated FRET value calculated from a Cy3-Cy5 Förster radius of 57 angstroms.

Single-molecule measurements were made using a solution confocal fluorescence microscope, in which FRET values are extracted from individual freely diffusing molecules as they traverse through the excitation beam. FRET values were collected for several thousand molecules under each condition and then compiled into FRET population histograms. The CR4/5 domain in the absence of TERT revealed a substantially heterogeneous FRET profile with values ranging across the FRET scale from ~0.1 to ~1, with the majority of molecules falling into populations centered at FRET~0.75 and ~0.9 (Fig. 6B). This observation was consistent with our chemical probing experiments with the same RNA fragment, which sampled an ensemble of distinct structural states. Molecules reporting high FRET values likely exist in a conformation in which the P6.1 stem loop is in close proximity to P6b, while lower FRET states indicated conformations of CR4/5, in which P6.1 is separated from P6b. Next, we measured the FRET properties of the dye-labeled CR4/5 domain after reconstitution with TERT and the hTR template/pseudoknot (t/PK) domain into catalytically active telomerase RNP complexes. Assembly into such telomerase RNPs essentially abolished the apparent heterogeneity of the CR4/5 domain with a single major FRET population centered around ~0.3 (Fig. 6C). This finding suggested that upon telomerase assembly the P6.1 and P6b stems are stabilized at an increased distance when compared to the major populations in a TERT-unbound structural ensemble. The estimated distance (~65-70 angstroms) between the FRET dyes in an assembled state is consistent with the respective dye label positions modeled in the human telomerase cryo EM structure (Fig. 6D) (Nguyen et al. 2018). This result lends additional support to a human CR4/5 structural transition upon binding to the TERT protein as was proposed for the medaka CR4/5 domain (Huang et al. 2014; Kim et al. 2014).

## DISCUSSION

Telomerase RNPs derived from diverse organisms must assemble upon highly structured telomerase RNA (TR) scaffolds (Zappulla and Cech 2006; Egan and Collins 2012a). TRs possess a multi-domain architecture conserved from unicellular ciliates to humans and serve to nucleate the assembly of telomerase complexes through interactions with the telomerase reverse transcriptase (TERT) and other lineage-specific proteins (Romero and Blackburn 1991; Chen et al. 2000; Chen and Greider 2004; Egan and Collins 2010). Despite their essential role in telomerase assembly, it remains unclear how TRs transition from their initial protein-free conformations to the intricate tertiary structures seen in active telomerase complexes (Jiang et al. 2018; Nguyen et al. 2018). In the present study, we demonstrate that the essential P6.1 stem of hTR CR4/5 is not stably folded in vitro and exists as a structural ensemble that is remodeled by the binding of TERT. Stabilizing P6.1 with engineered mutations in the 3WJ decreased the efficiency of endogenous telomerase assembly. Based upon these results, we propose a working model for telomerase RNP biogenesis that involves dynamic remodeling of a heterogeneous ensemble of hTR CR4/5 structural states (Fig. 7).

**Figure 7:**
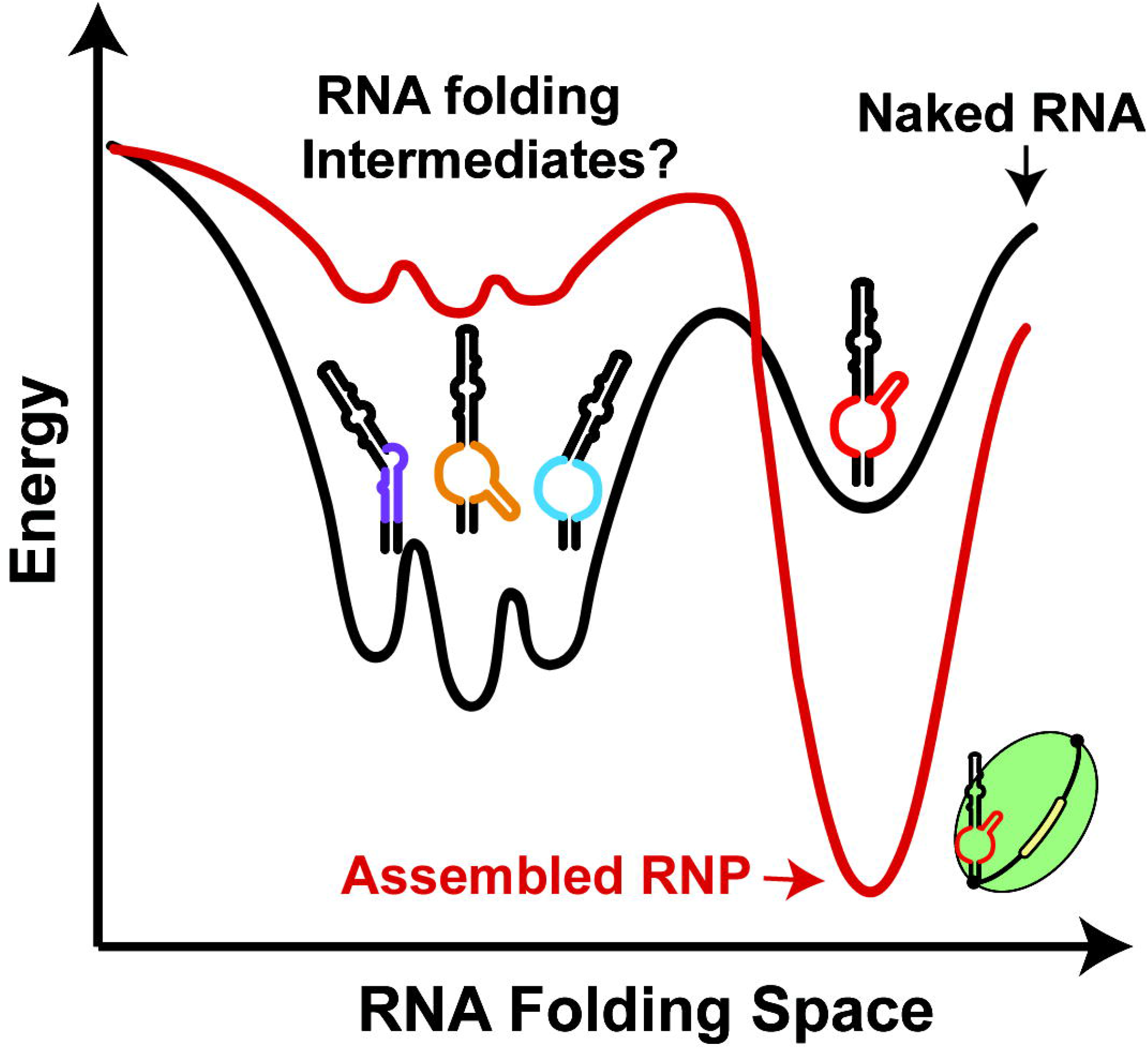
Model describing functional role of CR4/5 folding heterogeneity in human telomerase biogenesis. A schematic depicting a hypothetical folding landscape of the hTR CR4/5 domain. Energy valleys represent unique conformations available to CR4/5. The ‘depth’ of a valley is a conceptual proxy for the stability and relative abundance of a particular RNA conformation. In the folding landscape of the ‘naked’ CR4/5 RNA (black line), there exists a diverse ensemble of 3WJ conformations with a small contingent of molecules adopting a fold representative of the canonical P6.1 stem (red 3WJ). Upon RNP assembly, the CR4/5 folding landscape becomes dominated by one predominant CR4/5 conformation (red line), due to TERT-induced remodeling of the CR4/5 structure.

A protein-induced RNA structural rearrangement during telomerase assembly is expected to be essential at the stem terminal element (STE, stem-loop IV in *Tetrahymena* TR and CR4/5 in hTR). This region of TR makes a high affinity interaction with TERT (Bley et al. 2011) and when mutated, abrogates telomerase biogenesis (Mitchell and Collins 2000; Chen et al. 2002), precipitating human disease. In the *Tetrahymena* telomerase RNP, the TR stem-loop IV binds the assembly factor p65, which stabilizes a bent-helix conformation that places the apical loop at the interface of the TRBD and CTE domains of *Tetrahymena* TERT, potentially stabilizing the architecture of TERT (O’Connor and Collins 2006; Stone et al. 2007; Akiyama et al. 2012; Singh et al. 2012; Jiang et al. 2018). In hTR, the H/ACA box proteins (Dyskerin, NOP10, NHP2, and GAR1) regulate telomerase biogenesis and may play a similar role in facilitating the CR4/5 to adopt a conformation that engages the TRBD-CTE interface (Egan and Collins 2012b; Chen et al. 2018). Structural studies of the smaller *Oryzias latipes* (medaka) CR4/5 revealed protein-induced rearrangements of the 3WJ motif, rotating the P6.1 stem nearly 180 degrees around the axis of P5 and P6 to clamp upon the TERT RNA binding domain (TRBD) (Huang et al. 2014; Kim et al. 2014). Presumably, the hTR CR4/5 adopts a similar RNP assembled conformation given it shares invariant nucleotides comprising the P6.1 region and most of the 3WJ motif, a notion consistent with the medium-resolution cryoEM structure of human telomerase (Nguyen et al. 2018). However, the human 3WJ is expanded by ten nucleotides compared to its medaka counterpart and therefore might traverse a more complex folding landscape to arrive at its functional RNP state.

The observation of high 1M7-reactivity in nucleotides proposed to form the essential P6.1 stem of hTR CR4/5 was initially surprising, given that 1M7 does not react with the comparable nucleotides in the medaka CR4/5 (Fig. 3). Using these data to guide *in silico* prediction of medaka CR4/5 structure recovered the secondary structure consistent with its atomic resolution model and with high statistical confidence by bootstrap subsampling (Fig. 2 and 3). In contrast, including SHAPE data in the prediction of hTR CR4/5 lowered the statistical confidence of P6.1 (Fig. 2 and 3). Manual inspection of predicted secondary structures resulting from bootstrapping analysis revealed many hTR CR4/5 conformations with alternate base pairing configurations, suggesting the 3WJ and P6.1 stem of hTR CR4/5 can be structurally heterogeneous. An exhaustive mutate-and-map strategy (Kladwang et al. 2011a; Tian et al. 2014) of hTR CR4/5 identified base pairing signatures between specific nucleotides in P5 and P6, but was unable to detect Watson-Crick base pairing between nucleotides proposed to form P6.1 (Fig. 4). Notably, two G->C substitutions, G61C and G63C (G303C and G305C in full length hTR) abolished the 1M7 reactivity of the P6.1 region. RNAstructure predictions of these mutants reveal a non-3WJ conformation, in which nucleotides from the P6.1 region pair with nucleotides from P5 (Fig. S6). Interestingly, a patient derived hTR mutation at position G305 disrupts RNP assembly and induces aplastic anemia (Yamaguchi et al. 2003). These data suggest that bases in the P6.1 region may serve as ‘linchpins’, serving to stabilize the overall folding of the 3WJ motif.

The sequence of the P6.1 stem and flanking regions are strictly conserved across vertebrate TRs. Covariance patterns in CR4/5 suggest evolutionary pressure to maintain the P5 and P6 stems (Chen and Greider 2004), whereas the P6.1 stem lacks any instances of co-varying base pairs. Yet, it is known that a stable P6.1 stem is required for TERT binding (Mitchell and Collins 2000; Chen et al. 2002; Bley et al. 2011; Kim et al. 2014). The extreme sequence conservation within P6.1 stem loop and junction region of hTR suggests the presence of a selective pressure other than preservation of RNA structure alone. One explanation for these seemingly discordant findings could be that the P6.1 stem is characteristic of the functionally ‘assembled’ state of hTR CR4/5, but in the absence of TERT the domain adopts conformations other than the canonical P6.1 stem (Fig. 7). This conclusion is directly supported by the results of our smFRET analysis, in which we observe a heterogeneous FRET distribution for hTR CR4/5 only in the absence of TERT. Moreover, hTR sequence variants engineered to stabilize the P6.1 stem elicit telomerase RNP assembly defects to varying degrees in cell transfection assays. Taken together, these results suggest the presence of alternate conformations within the hTR expanded junction region, which ultimately must adopt the essential P6.1 stem loop during telomerase RNP biogenesis. The specific identities and functional role(s) of such putative hTR folding intermediates remain to be determined; however, it is conceivable that junction nucleotides may be conserved to preserve RNA structural plasticity required for RNP assembly, as well as to mediate sequence-specific protein interactions that may or may not be present in the fully assembled RNP complex. Lastly, recent reports have suggested additional non-canonical roles of hTR that may also explain the patterns of RNA sequence conservation in CR4/5 junction region (Rubtsova et al. 2018; Laudadio et al. 2019).

The assembly of large ribonucleoprotein complexes is a complex cellular task. The ability to modulate the conformation of an RNA scaffold may represent a point of cellular control over when and how RNP biogenesis occurs. Indeed, telomerase assembly from hTR and TERT is a point of regulation during telomere maintenance (Kellermann et al. 2015) and improper telomerase assembly is implicated in telomere syndromes (Mitchell and Collins 2000). Here, we have characterized structural heterogeneity in a catalytically essential region of hTR, and present evidence that RNA conformational intermediates may play a functional role in telomerase assembly. As the potential of RNA as a therapeutic target continues to grow (Dowdy 2017), we consider hTR biogenesis intermediates promising targets for the development of new telomerase inhibitors.

## MATERIALS AND METHODS

### Preparation of RNAs for chemical probing and *in vitro* telomerase reconstitution

#### Design and synthesis of RNA chemical probing constructs

Constructs for RNA chemical probing contained the RNA of interest (medaka CR4/5 (nt 170-220) and hTR CR4/5 (nt 243-326)) with additional flanking sequences for normalization purposes in data analysis (described below) and for reverse-transcriptase binding (Kladwang et al. 2014). RNA constructs for chemical probing were iteratively queried on the RNAstructure webserver (Reuter and Mathews 2010) and re-designed to discourage base pairing of the flanking sequences with the RNA of interest. Each RNA construct was synthesized by in vitro transcription. The DNA templates were assembled from DNA oligonucleotides designed using the Primerize tool (Tian et al. 2015) and synthesized by IDT (Fig. S7 and Table S2). In the event that a complete DNA template could not be synthesized by one primer assembly reaction using Phusion polymerase (NEB), a ‘two-piece’ scheme was employed, in which the products of two separate primer assemblies were used to generate the complete DNA product.

#### In vitro transcription of RNAs

RNA constructs for chemical probing and fragments used for *in vitro* telomerase reconstitution (hTR CR4/5 (nt 239-328) and hTR t/PK (nt 32-195)) were in vitro transcribed using homemade T7 RNA polymerase (Rio 2013) in RNA polymerase reaction buffer (40 mM Tris-HCl, pH 7.9, 28 mM MgCl_2_, 90 mM DTT, 2 mM spermidine, 1.5 mM each NTP and 40 U RNasin Plus (Promega)). The reaction was incubated overnight at 37°C followed by the addition of 10 units of TURBO DNase (Thermo Fisher) for 15 min at 37°C. RNA was phenol-chloroform extracted and ethanol precipitated prior to denaturing urea polyacrylamide gel electrophoresis (PAGE) purification. RNAs used in mutate-and-map experiments were transcribed in parallel on 96-well plates and purified using AMPure XP beads (Agencourt). RNA quality was then checked diagnostically by denaturing urea PAGE.

### Structural modeling of RNAs guided by chemical probing data

#### Chemical probing of RNAs

Chemical probing and mutate-and-map experiments were carried out as described previously (Kladwang and Das 2010; Cordero et al. 2014; Kladwang et al. 2014). Briefly, 1.2 pmol of RNA was denaturated at 95°C in 50 mM Na-HEPES, pH 8.0, for 3 min, and folded by cooling to room temperature over 20 min, and adding MgCl_2_ to 10 mM. RNA was aliquoted in 15 μl volumes into a 96-well plate and mixed with nuclease-free H_2_O (control), or chemically modified in the presence of 5 mM 1-methyl-7-nitroisatoic anhydride (1M7, provided by Dr. Manny Ares, UCSC), 25 mM 1-cyclohexyl-(2-morpholinoethyl) carbodiimide metho-p-toluene sulfonate (CMCT, Sigma Aldrich), or 0.25% dimethyl sulfate (DMS, Sigma Aldrich) for 10 min at room temperature. Mutate-and-map experiments utilized only 1M7 as the chemical modifier. Chemical modification was stopped by adding 9.75 μl quench and purification mix (1.53 M NaCl, 1.5 μl washed oligo-dT beads (Ambion), 6.4 nM FAM-labeled reverse-transcriptase primer (sequence in Supplementary Table 1), and 2.55 M Na-MES for 1M7 and CMCT reactions, or 50% 2-mercaptoethanol for DMS reactions. RNA in each well was purified by bead immobilization on a magnetic rack and two washes with 100 μL 70% ethanol. RNA was then resuspended in 2.5 μl nuclease-free water prior to reverse transcription.

#### Reverse transcription of modified RNAs and cDNA purification

RNA was reverse-transcribed from annealed fluorescent primer in a reaction containing 1x First Strand Buffer (Thermo Fisher), 5 mM DTT, 0.8 mM dNTP mix, and 20 U of SuperScript III Reverse Transcriptase (Thermo Fisher) at 48°C for 30 mins. RNA was hydrolyzed in the presence of 200 mM NaOH at 95°C for 3 min, then placed on ice for 3 min and quenched with 1 volume 5 M NaCl, 1 volume 2 M HCl, and 1 volume 3 M sodium acetate. cDNA was purified on magnetic beads as described previously, then eluted by incubation for 20 min in 11 μl Formamide-ROX 350 mix (1000 μl Hi-Di Formamide (Thermo Fisher) and 8 μL ROX 350 ladder (Thermo Fisher)). Samples were then transferred to a 96-well plate in ‘concentrated’ (4 μl sample + 11 μl ROX mix) and ‘dilute’ (1 μl sample + 14 μl ROX mix) for saturation correction in downstream analysis. Sample plates were sent to Elim Biopharmaceuticals for analysis by capillary electrophoresis.

#### Analysis of capillary electrophoresis data with HiTRACE

Capillary electrophoresis runs from chemical probing and mutate-and-map experiments were analyzed with the HiTRACE MATLAB package (Yoon et al. 2011). Lanes of similar treatment groups (e.g. 1M7 modified) were aligned together, bands fit to Gaussian peaks, background subtracted using the no-modification lane, corrected for signal attenuation and normalized to the internal hairpin control. The end result of these steps is a numerical array of ‘reactivity’ values for each RNA nucleotide that can be used as weights in structure prediction. For mutate-and-map datasets, each nucleotide is assigned a Z-score, calculated as its average reactivity across all mutants divided by the standard deviation (Kladwang et al. 2011a). Nucleotides with overall high reactivity across the mutants (average of 0.8 or higher) are ignored in Z-score calculation.

#### Data-guided RNA structure prediction

Data-guided secondary structure modeling was performed using the Biers MATLAB package (https://ribokit.github.io/Biers/). Briefly, the Fold function of the RNAstructure suite applied reactivity values as pseudoenergy modifiers to calculate the minimum free energy structure of CR4/5 RNA. Bootstrapping analysis of data-guided structure prediction was performed as described previously (Kladwang et al. 2011b; Tian et al. 2014). Briefly, the reactivity array was subsampled with replacement one hundred times and used to calculate the minimum free energy RNA structure. This generated an artificial ensemble from which each unique helical element observed could be assigned a percentage value based on its abundance. These percentage values represent statistical confidence of a specific helix forming in a secondary structure model. For mutate-and-map datasets, Z-scores were used as pseudo-energy modifiers to calculate a base pairing probability matrix with RNAstructure and to run bootstrapping analysis with Biers. Secondary structures were visualized using the VARNA applet (Darty et al. 2009).

### Telomerase expression and purification

#### In vitro reconstitution of human telomerase

Human telomerase was reconstituted in rabbit reticulocyte lysate (RRL) using the TNT Quick Coupled Transcription/Translation system (Promega) as described previously (Weinrich et al. 1997; Jansson et al. 2019). In Lo-bind tubes (Eppendorf), 200 μl of TnT quick mix was combined with 5 μg of pNFLAG-hTERT plasmid as well as 1 μM of in vitro transcribed and unlabeled hTR t/PK and CR4/5 fragments. Less abundant dye-labeled CR4/5 was added at 0.1 μM. The reaction was incubated for 3 h at 30°C. 5 μl of 0.5 M EDTA, pH 8.0, were then added to chelate Mg^2+^ ions present in the lysate. Human telomerase was immunopurified via the N-terminal FLAG tag on hTERT using αFLAG M2-agarose beads (Sigma-Aldrich). Beads contained in 50 μl bead slurry were first washed three times with wash buffer (50 mM Tris-HCl, pH 8.3, 3 mM MgCl_2_, 2 mM DTT, 100 mM NaCl) with 30 sec centrifugation steps at 2350 rcf at 4°C after each wash. The beads were then blocked twice in blocking buffer (50 mM Tris-HCl, pH 8.3, 3 mM MgCl_2_, 2 mM DTT, 500 μg/ml BSA, 50 μg/ml glycogen, 100 μg/ml yeast tRNA) for 15 min under gentle agitation at 4°C followed by 30 sec centrifugation at 2350 rcf and removal of the supernatant. After blocking, the beads were resuspended in 200 μl blocking buffer and added to the telomerase reconstitution reaction in RRL. The beads and lysate were incubated for 2 h at 4°C under gentle agitation. The beads were then pelleted for 30 sec at 2350 rcf and at 4°C and the supernatant was discarded. The beads were then washed three times in wash buffer containing 300 mM NaCl followed by three wash steps in wash buffer containing 100 mM NaCl. A 30 sec centrifugation at 2350 rcf at 4°C was performed between each wash cycle. To elute the enzyme, the beads were incubated in 60 μl elution buffer (50 mM Tris-HCl, pH 8.3, 3 mM MgCl_2_, 2 mM DTT, 750 μg/ml 3xFLAG peptide, 20% glycerol) under gentle agitation at 4°C for 1 h. After elution, the beads were removed by centrifugation at 10000 rcf through Nanosep MF 0.45 μm filters. 5 μl aliquots were prepared in Lo-bind tubes (Eppendorf), flash frozen in liquid nitrogen and stored at −80°C until use.

#### Reconstitution of human telomerase in HEK293T cells

For human telomerase expression in HEK293T cells, hTR and hTERT were overexpressed by transient transfection of the pBSU3hTR500 and pNFLAG-hTERT plasmids, respectively (Drosopoulos et al. 2005; Deshpande and Collins 2018). The hTR mutants (Mut-1 and Mut-2) were synthesized by a primer assembly reaction (Tian et al. 2015) and cloned into the pBSU3hTR500 vector. Plasmids were transfected using lipofectamine (ThermoFisher) in a T25 flask seeding the day prior with 2.5×10^6^ cells. Cells were transferred to a T75 flask the next day and allowed to culture overnight prior to harvesting. Cells were lysed with 500 μl ice cold CHAPS buffer (150 mM KCl, 50 mM Na-HEPES, pH 7.4, 0.1% CHAPS) and 3 μl RNAsin Plus. Cell debris was pelleted by centrifugation at >13,000 rpm at 4°C for 10 min. 40 μl aliquots of cleared lysate were flash frozen in liquid nitrogen for western blotting, while the rest of the lysate was mixed with 200 μl of a 1:1 slurry of αFLAG M2-agarose beads. Immunopurification, subsequent washes, and elution were performed as described above. Aliquots of 10 μl were flash frozen in liquid nitrogen.

### Telomerase activity assays

#### ^32^P-end-labeling of DNA primers

50 pmol of DNA primer was labeled with gamma-^32^P ATP using T4 polynucleotide kinase (NEB) in 1x PNK buffer (70 mM Tris-HCl, pH 7.6, 10 mM MgCl_2_, 5 mM DTT) in 50 μl reaction volume. The reaction was incubated for 1 h at 37°C followed by heat inactivation of T4 PNK at 65°C for 20 min. Centrispin columns (Princeton Separations) were used to purify labeled primer.

#### Primer extension assays

Telomerase activity assays of *in vitro* reconstituted human telomerase were performed using 5 μl purified telomerase in a 15 μl reaction volume brought to 1x activity buffer concentrations (50 mM Tris-HCl, pH 8.3, 50 mM KCl, 1 mM MgCl_2_, 2 mM DTT, 50 nM ^32^P-end-labled primer, and 10 μM of each dATP, dTTP, and dGTP). Reactions were incubated for 90 min at 30°C and quenched with 200 μl 1x TES buffer (10 mM Tris-HCl, pH 7.5, 1 mM EDTA, 0.1% SDS). DNA products were then phenol-chloroform extracted and ethanol precipitated. DNA pellets were resuspended in 1x formamide gel loading buffer (50 mM Tris Base, 50 mM boric acid, 2 mM EDTA, 80% (v/v) formamide, 0.05% (w/v) each bromophenol blue and xylene cyanol) and resolved on a 12% denaturing urea PAGE gel. The gel was then dried and exposed to a storage phosphor screen (GE Healthcare) and scanned using a Typhoon scanner (GE Healthcare). Band intensities were quantified using SAFA and ImageJ (Das et al. 2005; Schneider et al. 2012). The ‘fraction left behind’ (FLB) for a given lane was calculated by summing each repeat addition processivity (RAP) band and all RAP bands below it divided by the total RAP band intensity counts for that lane. The natural logarithm of (1-FLB) was then plotted against repeat number and fitted by linear regression. The slope value of the linear fit was used to determine processivity R_1/2_ values from -ln(2)/slope (Latrick and Cech 2010). Total activity was calculated in ImageJ by taking the total intensity of each lane and normalizing to the wild-type lane.

#### Western blots

Immunopurified telomerase complexes were resolved on a 10% SDS PAGE gel and transferred to a PVDF membrane (Millipore) using a semi-dry blotter (Biorad) at 13 V for 1 h. After blocking with 5 % (w/v) milk in TBS-T (20 mM Tris-HCl, pH 7.5, 150 mM NaCl, 0.1% Tween-20), the membrane was incubated with FLAG M2-Peroxidase antibody (Sigma Aldrich) at a 1:1000 dilution in 1 % (w/v) milk in TBS-T for 18 h at 4°C. After three 10 min wash steps with TBS-T, the membrane was immersed in Pierce western blot detection solution (Thermo Fisher) and imaged in a Biorad GelDoc device.

#### RNA dot blot

5 μl of immunopurified human telomerase was diluted to 10 μl in formamide loading buffer (50 mM Tris Base, 50 mM boric acid, 2 mM EDTA, 80% (v/v) formamide, 0.05% (w/v) each bromophenol blue and xylene cyanol) and heated at 70°C for 5 min and placed on ice. The sample was pipetted in droplets onto a Hybond N+ membrane (GE Lifesciences) and allowed to air dry at room temperature for 1 h before cross-linking in a UV transilluminator at 254 nm for 5 min. The membrane was then blocked with 50 ml Church buffer (1% BSA, 1 mM EDTA, 500 mM sodium phosphate, pH 7.2, 7% SDS) at 55°C for 1 h). Next, 3×10^6^ cpm of 5’ gamma-^32^P labeled DNA oligonucleotide probe (sequence: 5’-TATCAGCACTAGATTTTTGGGGTTGAATG- 3’) was added to the membrane and incubated under shaking at 55°C for at least 12 h. The membrane was washed three times with 0.1x saline-sodium-citrate buffer (15 mM NaCl, 1.5 mM trisodium citrate, pH 7.0, 0.1% SDS) while shaking for 15 min at room temperature. The membrane was imaged using a storage phosphor screen (GE Healthcare) and a typhoon scanner (GE Healthcare). Quantification of the signal was performed in ImageJ. To determine concentrations, samples were compared against in vitro transcribed telomerase RNA standards included in each blot.

### Preparation of dye-labeled hTR CR4/5 for single-molecule experiments

#### Synthesis of dye-labeled hTR CR4/5 RNA

Synthetic CR4/5 (hTR 239-330) was ordered from Dharmacon as two separate oligonucleotides: Fragment 1 (hTR 239-278) and Fragment 2 (hTR 279-330), each harboring a site-specific amino-allyl modification at the 5 position of uracil base as indicated in Supplementary Table 1. Oligonucleotides were de-protected in deprotection buffer (100 mM acetic acid, pH 3.6) following the manufacturer’s instructions, then ethanol precipitated in the presence of 300 mM sodium acetate, pH 5.2. To enable RNA ligation, Fragment 2 was phosphorylated using T4 PNK (NEB), phenol-chloroform extracted, and ethanol precipitated in presence of sodium acetate. 10 nmol of each RNA fragment were brought to 100 μl in 0.1 M sodium bicarbonate, pH 9.0, and mixed with an equal volume of an Cy3 or Cy5 Amersham mono-reactive dye pack in DMSO (GE Healthcare). The labeling mix was incubated at 37°C in the dark for 2 hours, then ethanol precipitated. Pellets were resuspended in 60 μl Buffer A (0.1 M triethylammonium acetate (TEAA), pH 7.5), and HPLC purified on a reversed phase C8 column (Agilent Technologies).

#### Ligation of synthetic RNA fragments

To generate a CR4/5 RNA (hTR 239-328) with fluorescent dyes at positions U274 and U312, a splinted ligation reaction (Akiyama and Stone 2009) containing 800 pmol of Cy3-labeled Fragment 2 (hTR 279-330), 1600 pmol of Cy5-labeled Fragment 1 (hTR 239-278), 1600 pmol of DNA splint (sequence: 5’-AGTGGGTGCCTCCGGAGAAGCCCCGGGCCGAC-3’) in 0.5x T4 DNA ligase buffer (NEB) was brought to 100 μl volume and incubated at 95°C for 5 min and at 30°C for 10 min. 100 μl ligation mix (1.5x T4 DNA ligase buffer, 4000 U T4 DNA ligase (NEB), 2 mM ATP and 1 U/ μl RNAsin Plus (Promega)) was added to the reaction and incubated at 30°C for 18 h. 10 U of TURBO DNAse (Thermo Fisher Scientific) were added and the reaction incubated at 37°C for 15 min. The RNA was phenol-chloroform extracted and ethanol precipitated prior to PAGE purification.

### Single-molecule experiments

#### Slide preparation for imaging

Glass micro slides (Gold Seal) were washed by hand with Alconox detergent and warm water, then dried with nitrogen. Sample channels were constructed with Parafilm strips and a plasma-cleaned glass coverslip (Fisher Scientific). Channels were blocked with 10 mg/ml BSA (NEB) for 1 h and washed with imaging buffer (50 mM Tris-HCl, pH 8.3, 50 mM KCl, 1 mM MgCl_2_, 1 mg/ml BSA, 8% glucose, and (±)-6-Hydroxy-2,5,7,8-tetramethylchromane-2-carboxylic acid (Trolox) at saturation. Trolox-containing imaging buffer was generally filtered (0.2 μm) before and after adjusting the pH to 8.3 with NaOH. For imaging, 0.01 volumes of ‘Gloxy’ solution (10 mM Tris-HCl, pH 8.0, 50 mM NaCl, 200 μg/mL catalase, 100 mg/ml glucose oxidase) were added to the imaging buffer.

#### Confocal microscopy of doubly labeled CR4/5 RNA and human telomerase

Data was acquired with a confocal fluorescence microscope with 200 pM labeled hTR CR4/5 and 50-fold diluted aliquots of *in vitro* reconstituted labeled human telomerase. A green laser (532 nm) set to 100 μW was used to excite the Cy3 donor dye within the slide channel, and fluorescence from a ~100 nm^3^ volume was collected through a pinhole and passed on to a dichroic mirror to separate green and red wavelengths. Red and green light were individually detected by avalanche photodiode detectors (APDs) and written to a data file using custom LabView software. Data was collected for 30 min, usually capturing fluorescence from thousands of individual molecules.

#### Analysis of single-molecule data

Using custom MATLAB scripts, the data was thresholded to include only molecules with Cy5 fluorescence one standard deviation above the mean intensity detected by the red (637 nm) APD, as well as corrected for direct Cy5 excitation by green light and dichroic mirror breakthrough. FRET efficiency was calculated in MATLAB with the equation FRET = I_A_/(I_A_+I_D_), where I_A_ and I_D_ are acceptor and donor intensity, respectively). Histograms were generated using GraphPad Prism. Gaussian approximation of FRET populations was performed by fitting each histogram with a nonlinear regression model, in which the mean of each Gaussian function was constrained to values determined by visual approximation.

## ACKNOWLEDGEMENTS

We thank Dr. Manuel Ares for the generous gift of 1M7 reagent made in his laboratory. We also thank Ann Kladwang and Joseph Yesselman for their guidance in setting up the chemical probing experiments. We thank Dr. Julian Chen and Dr. Kathleen Collins for the gifts of the hTR and hTERT transfection plasmids. The work was supported by an NIH R01 GM095850 to M.D.S and R35 GM122579 grant to R.D. J.H. was supported by an Swiss National Science Foundation Early Postdoc. Mobility Fellowship (P2EZP3_181605), and N.M.F. was supported on an NIH T32 GM133391.

