## Supplementary Information for "Folding heterogeneity in the essential human telomerase RNA three-way junction"

*Running title (50 characters):* Function of telomerase RNA structural heterogeneity

Christina Palka[1], Nicholas M. Forino[2], Jendrik Hentschel[1], Rhiju Das[3,4,5], Michael D. Stone[1,6]

1. Department of Chemistry and Biochemistry, University of California, Santa Cruz, California 95064, USA.
2. Department of Molecular, Cell, and Developmental Biology, University of California, Santa Cruz, California 95064, USA.
3. Biophysics Program, Stanford University, Stanford, California 94305, USA.
4. Department of Biochemistry, Stanford University, Stanford, California 94305, USA.
5. Department of Physics, Stanford University, Stanford, California 94305, USA.
6. Center for Molecular Biology of RNA, University of California, Santa Cruz, California 95064, USA.

### Supplementary Figure S1

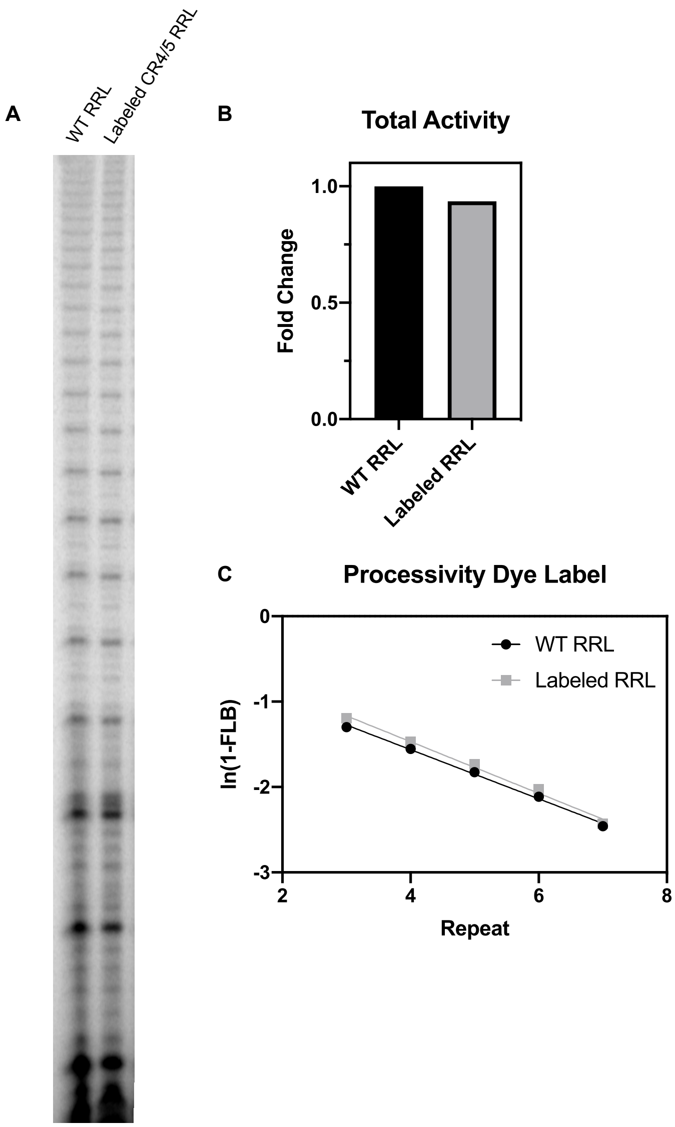


**Supplementary Figure S1: Primer extension assay of telomerase reconstituted with WT CR4/5 and dye-labeled CR4/5.** (**A**) Gel illustrating activity and processivity of reconstituted telomerase with WT CR4/5 and dye-labeled CR4/5 (**B**) Quantification of total activity. (**C**) Quantification of repeat addition processivity for telomerase enzymes reconstituted with unlabeled and dye-labeled CR4/5 domain (see Materials and Methods for reconstitution and data analysis details).

### Supplementary Figure S2

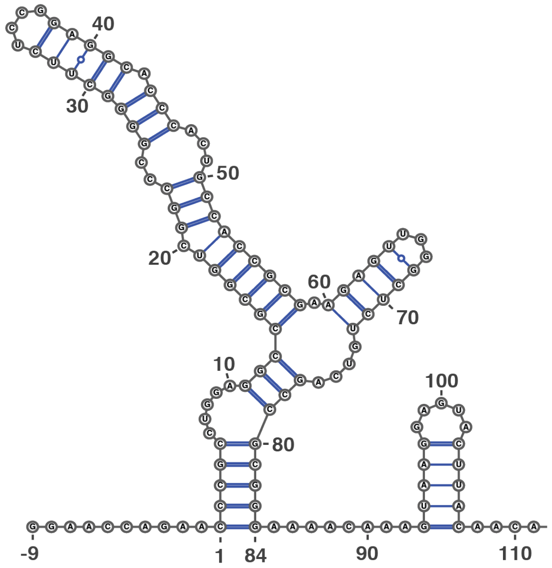


**RNAstructure Energy = -48.8**

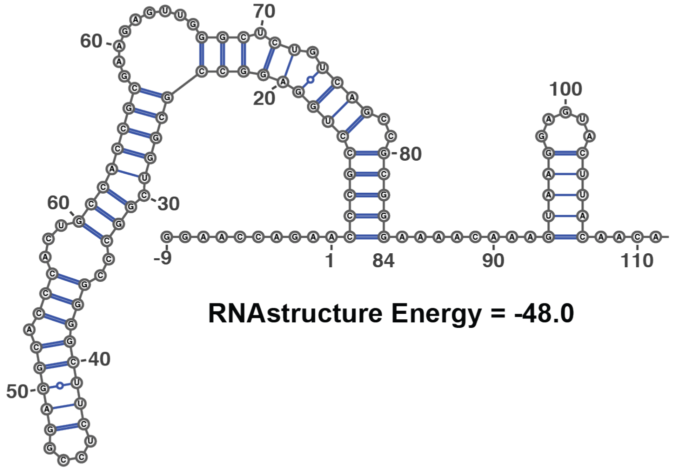


**RNAstructure Energy = -48.0**

**Supplementary Figure S2: Comparison of structures of the human CR4/5 domain predicted using the RNAstructure software package.** Similar energy values are observed for different structures predicted for the human CR4/5 domain.

### Supplementary Figure S3

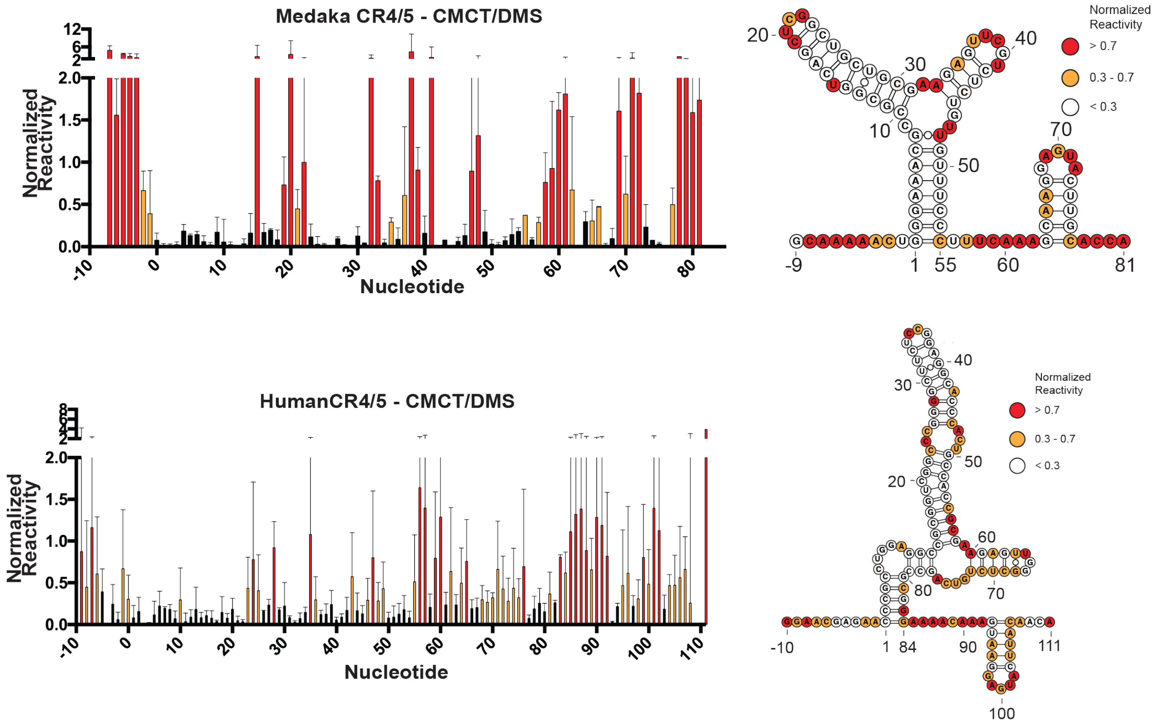


**Supplementary Figure S3: Chemical mapping of medaka and human CR4/5 domains using merged CMCT/DMS data.** Chemical mapping of the medaka (**A**) and human (**B**) CR4/5 domain using merged CMCT/DMS data. Plotted normalized reactivity values are color-coded (red > 0.7, yellow 0.3 - 0.7, and white < 0.3). Bar plot represents experiments conducted in triplicate and error bars are the standard deviation of the three replicates. Color-coded schematic of the reactivity data is shown on the RNAstructure predicted secondary structure.

### Supplementary Figure S4

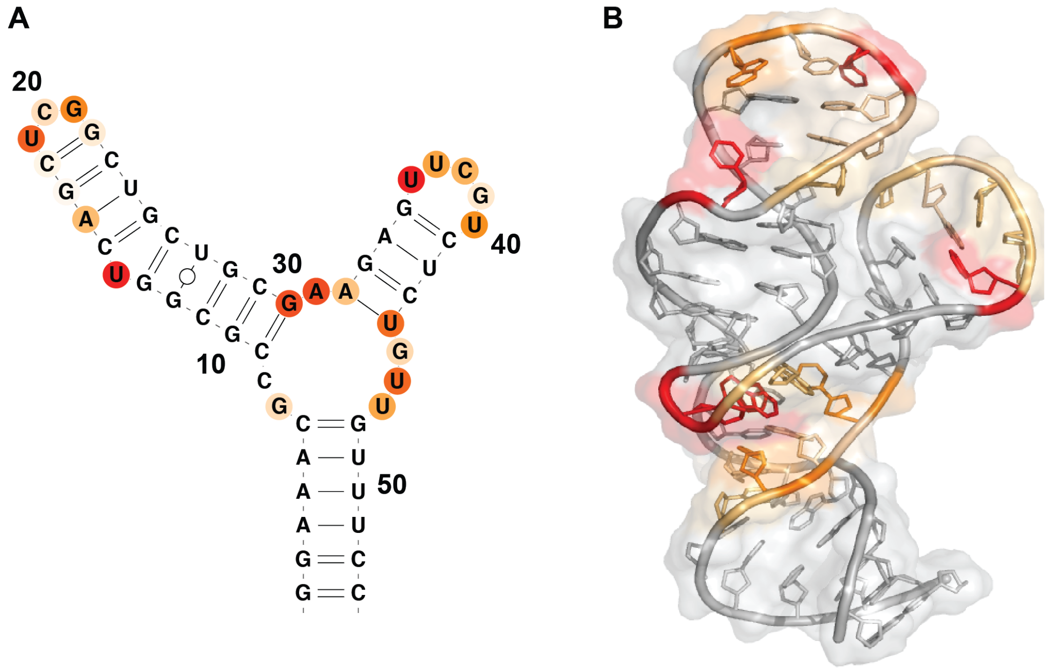


**Supplememtary Figure S4: Chemical mapping of medaka 1M7 mapped onto predicted secondary structure (A) and NMR solution structure (B) (PDB: 2MHI).**

### Supplementary Figure S5

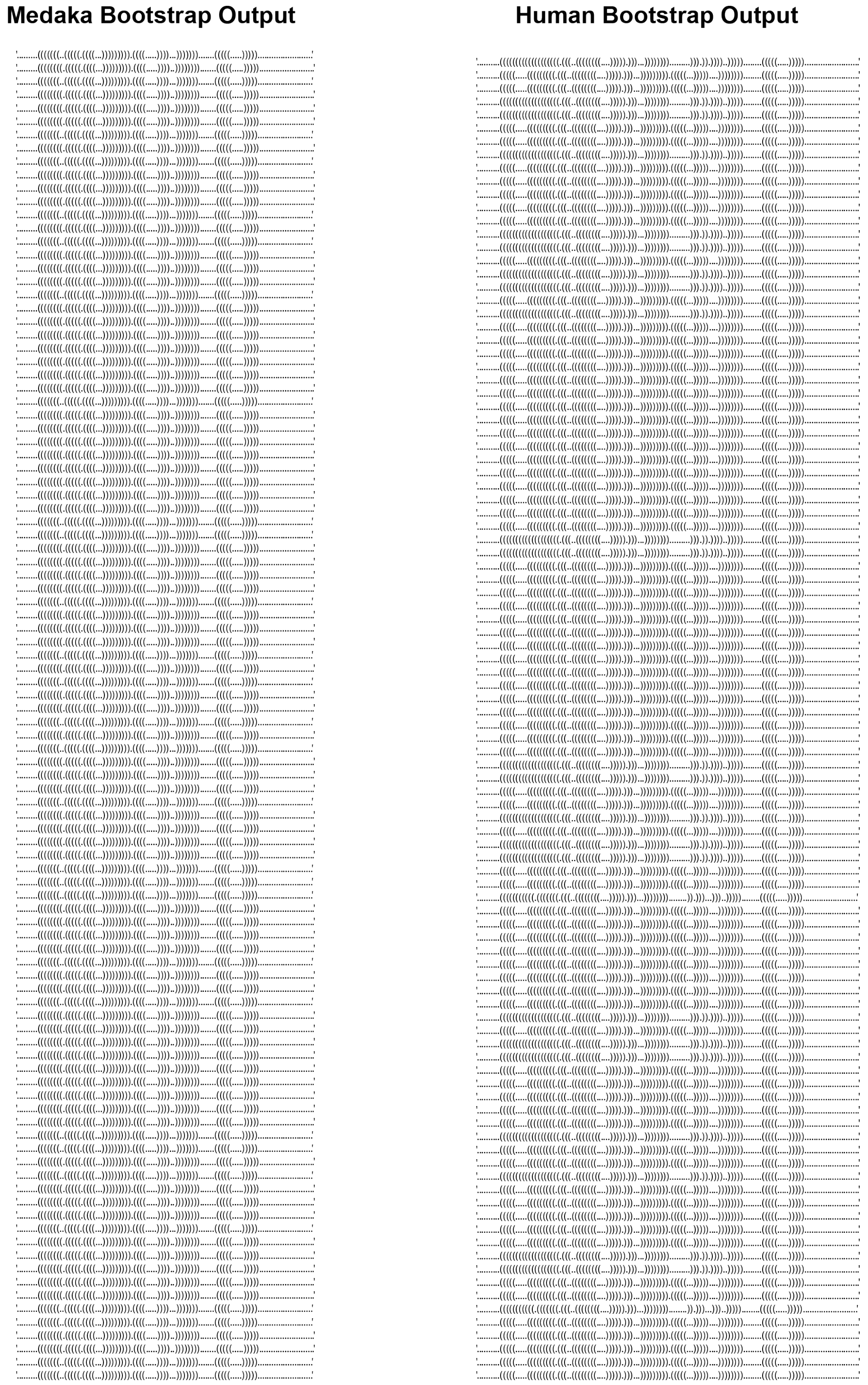


**Supplementary Figure S5: Comparison of dot-bracket structure output from bootstrapped structure prediction of the medaka and human CR4/5 domain.** Data-driven structure prediction for human and medaka CR4/5 domains. The above structures were predicted after 100 prediction iterations using Biers. Structures are shown in dot-bracket notation with “.” denoting a nucleotide that is not base-paired, and “(” denoting one side of a base-pair and “)” the other side of the base pair.

**Supplementary Figure S6**

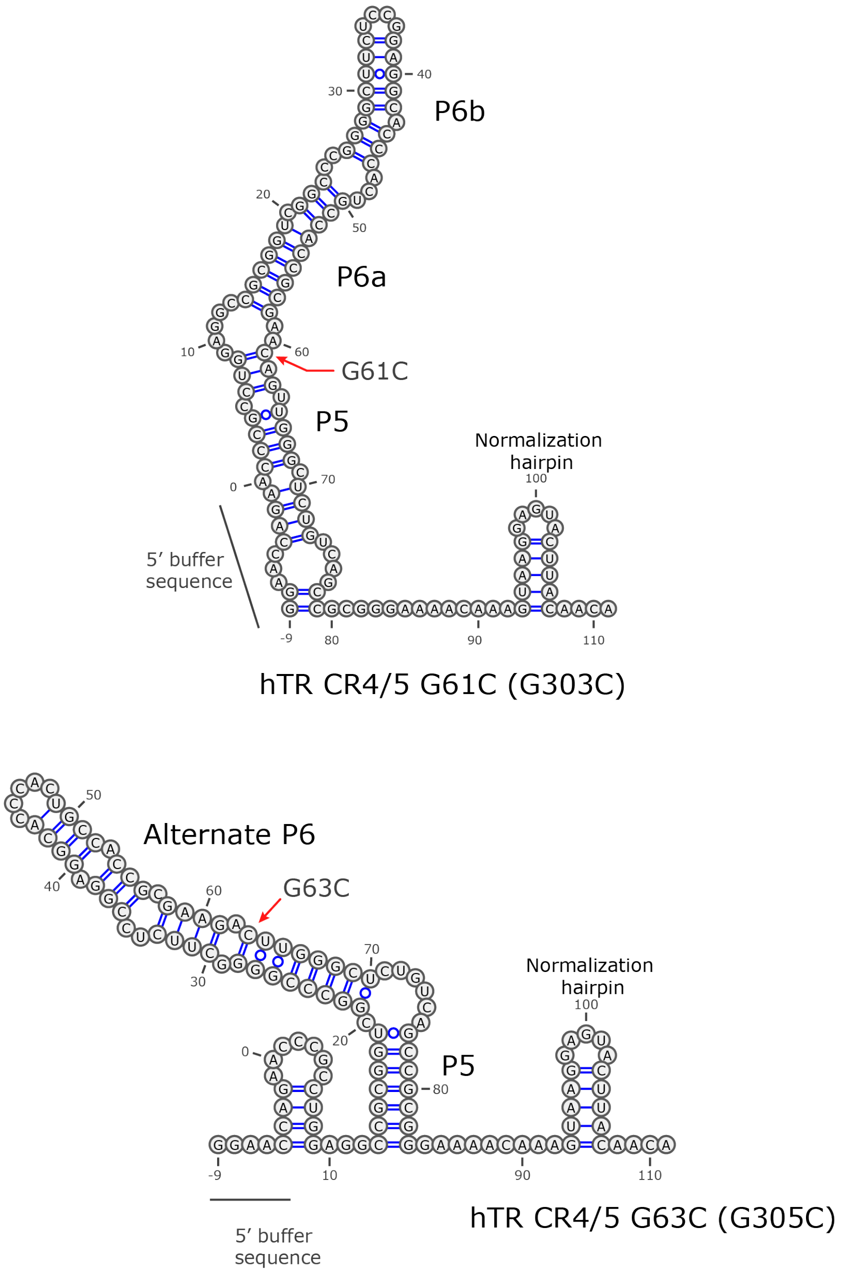


**Supplementary Figure S6: Data-driven structure prediction of select CR4/5 mutants G61C and G63.** The G61C mutation induced an alternate predicted conformation of the CR4/5 domain that resulted in the 5’ buffer annealing to the 3WJ junction region and part of P5 forming a base pairing interaction with the bases normally involved in the P6.1 stem-loop. The G63C mutant caused a dramatic rearrangement in the predicted structure resulting in a totally different structure, with the P6, P5 and 3WJ region all taking on new base-pairing partners. In both constructs the normalization hairpin formed correctly and the mutation made is indicated with a red arrow.

**Supplementary Figure S7**

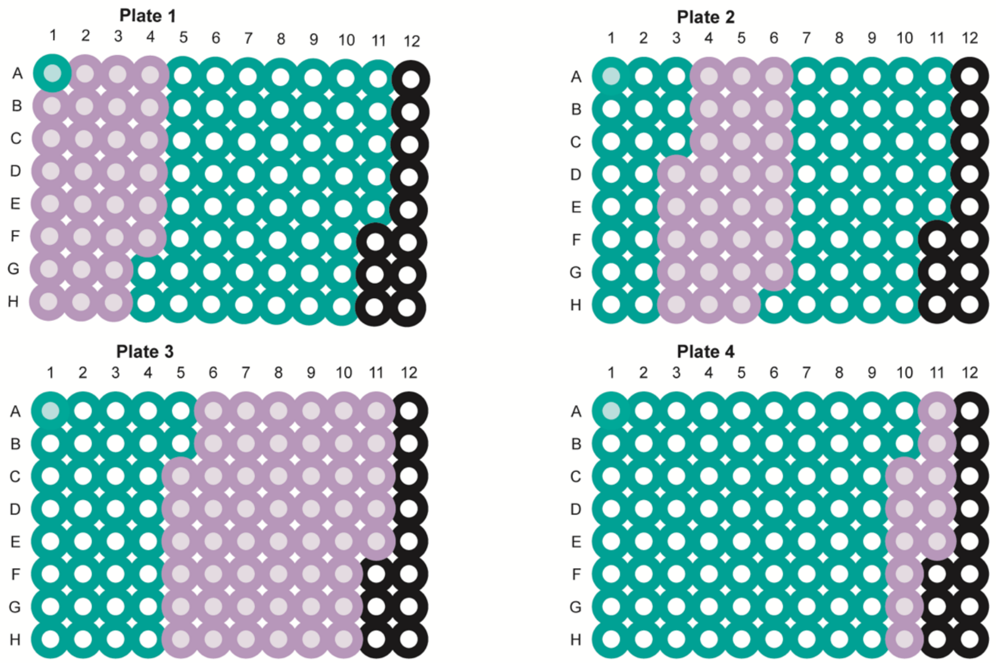


**Supplementary Figure S7: Plate layout for assembling the 84 Mutate-and-Map constructs.** Plate 1, 2, 3, and 4 are all illustrated. For each RNA four primers were used (one from each plate) for the in the primer assembly reaction. By matching the identities of the wells (A1, B1, etc) on each plate, a full-length mutated construct could be assembled (ie - using A1 primers from all plates will make WT CR4/5, using B1 primers from all plates will make mutant C1G, using C1 primers from all will make mutant C2G). Green corresponds to a WT primer for that plate and purple corresponds to a mutant. Only mutant primers (purple) are listed in the M2 oligo table and the primer listed as A1 on each plate lists the WT primer.

| Construct | Sequence |
|---|---|
| Human CR4/5 | TTCTAATACGACTCACTATAGGAACGAGAACCCGCCTG |
| Human CR4/5 | GGGTGCCTCCGGAGAAGCCCCGGGCCGACCGCGGCCTCCAGGCGGGTTCTCGTT |
| Human CR4/5 | TCCGGAGGCACCCACTGCCACCGCGAAGAGTTGGGCTCTGTCAGCCGCGGGA |
| Human CR4/5 | GTTGTTGTTGTTGTTTCTTTTGTTGTAAGTACTCCTTACTTTGTTTTCCCGCGGCTGA |
| Medaka CR4/5 | TTCTAATACGACTCACTATAGCAAAAACTGGGAAACGCCGCGGTCA |
| Medaka CR4/5 | AGGGAAACAACAGAGACGAACTCTTCGCAGCAGCCGAGCTGACCGCGGCGTT |
| Medaka CR4/5 | TCGTCTCTGTTGTTTCCCTTTCAAAGCAAGGAGTACTTGCACCAGACACAACACAAT |
| Medaka CR4/5 | GCTGTAAATTGTGTTGTGTCTGGTGCAAGT |
| Mut-1 (lock 7) | TTCTAATACGACTCACTATAGGAACGAGAACCCGCAAAAAAAACGCGGTCGGC |
| Mut-1 (lock 7) | GCGGTGGCAGTGGGTGCCTCCGGAGAAGCCCCGGGCCGACCGCGTT |
| Mut-1 (lock 7) | CCACTGCCACCGCGAAGAGTTGGGCTCTAAAAAAAGCGGGAAAACAAAGTAAGG |
| Mut-1 (lock 7) | GTTGTTGTTGTTGTTTCTTTTGTTGTAAGTACTCCTTACTTTGTTTTCCCGCTTTTTTT<br>A |
| Mut-2 (lock 3) | TTCTAATACGACTCACTATAGGAACCAGAACCCGCCTG |
| Mut-2 (lock 3) | GGGTGCCTCCGGAGAAGCCCCGGGCCGACCGCGGCCTCCAGGCGGGTTCTGGT |
| Mut-2 (lock 3) | TCCGGAGGCACCCACTGCCACCGCGACGAGCTGGGCTCGGTCAGCCGCGGGA |
| Mut-2 (lock 3) | GTTGTTGTTGTTGTTTCTTTTGTTGTAAGTACTCCTTACTTTGTTTTCCCGCGGCTGA |
| RT FAM primer (medaka) | /56-FAM/AAAAAAAAAAAAAAAAAAAAGCTGTAAATTGTGTTGTGTC |
| RT FAM primer (human) | /56-FAM/AAAAAAAAAAAAAAAAAAAAGTTGTTGTTGTTGTTTCTTT |

**Supplementary Table 1 - Oligonucleotides used in study** (except mutate-and-map)

| | |
|---|---|
| Telo primer for primer extension | TTAGGGTTAGGGTTAGGG |
| Dot Blot Hybridization Oligo | CGG TGG AAG GCG GCA GGC CGA GGC |
| hTR CR4/5 Fragment 1 | GAACCCCGCCUGGAGGCCGCGGUCGGCCCGGGGCU(5-LC-N-U)CUCC3 |
| hTR CR4/5 Fragment 2 | GGAGGCACCCACUGCCACCGCGAAGAGUUGGGC(5-LC-N-U)CUGUCAGCCGCGGGUCUC3 |
| DNA splint for CR4/5 construction | AGTGGGTGCCTCCGGAGAAGCCCCGGGCCGAC3 |

### Supplementary Table 2 - DNA primers used to make Mutate-and-Map templates

| PLATE 1 | | |
|---|---|---|
| **WellPosition** | **Name** | **Sequence** |
| *A01* | Lib1-WT | TTCTAATACGACTCACTATAGGAACGAGAACCCGCCTGGAGGCCGCGGTCGGCCCGGGG |
| *B01* | Lib1-C31G | TTCTAATACGACTCACTATAGGAACGAGAAGCCGCCTGGAGGCCGCGGTCGGCCCGGGG |
| *C01* | Lib1-C32G | TTCTAATACGACTCACTATAGGAACGAGAACGCGCCTGGAGGCCGCGGTCGGCCCGGGG |
| *D01* | Lib1-C33G | TTCTAATACGACTCACTATAGGAACGAGAACCGGCCTGGAGGCCGCGGTCGGCCCGGGG |
| *E01* | Lib1-G34C | TTCTAATACGACTCACTATAGGAACGAGAACCCCCCTGGAGGCCGCGGTCGGCCCGGGG |
| *F01* | Lib1-C35G | TTCTAATACGACTCACTATAGGAACGAGAACCCGGCTGGAGGCCGCGGTCGGCCCGGGG |
| *G01* | Lib1-C36G | TTCTAATACGACTCACTATAGGAACGAGAACCCGCGTGGAGGCCGCGGTCGGCCCGGGG |
| *H01* | Lib1-T37A | TTCTAATACGACTCACTATAGGAACGAGAACCCGCCAGGAGGCCGCGGTCGGCCCGGGG |
| *A02* | Lib1-G38C | TTCTAATACGACTCACTATAGGAACGAGAACCCGCCTCGAGGCCGCGGTCGGCCCGGGG |
| *B02* | Lib1-G39C | TTCTAATACGACTCACTATAGGAACGAGAACCCGCCTGCAGGCCGCGGTCGGCCCGGGG |
| *C02* | Lib1-A40T | TTCTAATACGACTCACTATAGGAACGAGAACCCGCCTGGTGGCCGCGGTCGGCCCGGGG |
| *D02* | Lib1-G41C | TTCTAATACGACTCACTATAGGAACGAGAACCCGCCTGGACGCCGCGGTCGGCCCGGGG |
| *E02* | Lib1-G42C | TTCTAATACGACTCACTATAGGAACGAGAACCCGCCTGGAGCCCGCGGTCGGCCCGGGG |
| *F02* | Lib1-C43G | TTCTAATACGACTCACTATAGGAACGAGAACCCGCCTGGAGGGCGCGGTCGGCCCGGGG |
| *G02* | Lib1-C44G | TTCTAATACGACTCACTATAGGAACGAGAACCCGCCTGGAGGCGGCGGTCGGCCCGGGG |
| *H02* | Lib1-G45C | TTCTAATACGACTCACTATAGGAACGAGAACCCGCCTGGAGGCCCCGGTCGGCCCGGGG |
| *A03* | Lib1-C46G | TTCTAATACGACTCACTATAGGAACGAGAACCCGCCTGGAGGCCGGGGTCGGCCCGGGG |
| *B03* | Lib1-G47C | TTCTAATACGACTCACTATAGGAACGAGAACCCGCCTGGAGGCCGCCGTCGGCCCGGGG |
| *C03* | Lib1-G48C | TTCTAATACGACTCACTATAGGAACGAGAACCCGCCTGGAGGCCGCGCTCGGCCCGGGG |
| *D03* | Lib1-T49A | TTCTAATACGACTCACTATAGGAACGAGAACCCGCCTGGAGGCCGCGGACGGCCCGGGG |
| *E03* | Lib1-C50G | TTCTAATACGACTCACTATAGGAACGAGAACCCGCCTGGAGGCCGCGGTGGGCCCGGGG |
| *F03* | Lib1-G51C | TTCTAATACGACTCACTATAGGAACGAGAACCCGCCTGGAGGCCGCGGTCCGCCCGGGG |
| *G03* | Lib1-G52C | TTCTAATACGACTCACTATAGGAACGAGAACCCGCCTGGAGGCCGCGGTCGCCCCGGGG |

| | | |
|---|---|---|
| *H03* | Lib1-C53G | TTCTAATACGACTCACTATAGGAACGAGAACCCGCCTGGAGGCCGCGGTCGGGCCGGGG |
| *A04* | Lib1-C54G | TTCTAATACGACTCACTATAGGAACGAGAACCCGCCTGGAGGCCGCGGTCGGCGCGGGG |
| *B04* | Lib1-C55G | TTCTAATACGACTCACTATAGGAACGAGAACCCGCCTGGAGGCCGCGGTCGGCCGGGGG |
| *C04* | Lib1-G56C | TTCTAATACGACTCACTATAGGAACGAGAACCCGCCTGGAGGCCGCGGTCGGCCCCGGG |
| *D04* | Lib1-G57C | TTCTAATACGACTCACTATAGGAACGAGAACCCGCCTGGAGGCCGCGGTCGGCCCGCGG |
| *E04* | Lib1-G58C | TTCTAATACGACTCACTATAGGAACGAGAACCCGCCTGGAGGCCGCGGTCGGCCCGGCG |
| *F04* | Lib1-G59C | TTCTAATACGACTCACTATAGGAACGAGAACCCGCCTGGAGGCCGCGGTCGGCCCGGGC |
| | **PLATE 2** | |
| **WellPosition** | **Name** | **Sequence** |
| *A01* | Lib1-WT | GGGTGCCTCCGGAGAAGCCCCGGGCCGA |
| *D03* | Lib1-T49A | GGGTGCCTCCGGAGAAGCCCCGGGCCGT |
| *E03* | Lib1-C50G | GGGTGCCTCCGGAGAAGCCCCGGGCCCA |
| *F03* | Lib1-G51C | GGGTGCCTCCGGAGAAGCCCCGGGCGGA |
| *G03* | Lib1-G52C | GGGTGCCTCCGGAGAAGCCCCGGGGCGA |
| *H03* | Lib1-C53G | GGGTGCCTCCGGAGAAGCCCCGGCCCGA |
| *A04* | Lib1-C54G | GGGTGCCTCCGGAGAAGCCCCGCGCCGA |
| *B04* | Lib1-C55G | GGGTGCCTCCGGAGAAGCCCCCGGCCGA |
| *C04* | Lib1-G56C | GGGTGCCTCCGGAGAAGCCCGGGGCCGA |
| *D04* | Lib1-G57C | GGGTGCCTCCGGAGAAGCCGCGGGCCGA |
| *E04* | Lib1-G58C | GGGTGCCTCCGGAGAAGCGCCGGGCCGA |
| *F04* | Lib1-G59C | GGGTGCCTCCGGAGAAGGCCCGGGCCGA |
| *G04* | Lib1-C60G | GGGTGCCTCCGGAGAACCCCCGGGCCGA |
| *H04* | Lib1-T61A | GGGTGCCTCCGGAGATGCCCCGGGCCGA |
| *A05* | Lib1-T62A | GGGTGCCTCCGGAGTAGCCCCGGGCCGA |
| *B05* | Lib1-C63G | GGGTGCCTCCGGACAAGCCCCGGGCCGA |
| *C05* | Lib1-T64A | GGGTGCCTCCGGTGAAGCCCCGGGCCGA |
| *D05* | Lib1-C65G | GGGTGCCTCCGCAGAAGCCCCGGGCCGA |
| *E05* | Lib1-C66G | GGGTGCCTCCCGAGAAGCCCCGGGCCGA |
| *F05* | Lib1-G67C | GGGTGCCTCGGGAGAAGCCCCGGGCCGA |
| *G05* | Lib1-G68C | GGGTGCCTGCGGAGAAGCCCCGGGCCGA |
| *H05* | Lib1-A69T | GGGTGCCACCGGAGAAGCCCCGGGCCGA |
| *A06* | Lib1-G70C | GGGTGCGTCCGGAGAAGCCCCGGGCCGA |
| *B06* | Lib1-G71C | GGGTGGCTCCGGAGAAGCCCCGGGCCGA |
| *C06* | Lib1-C72G | GGGTCCCTCCGGAGAAGCCCCGGGCCGA |
| *D06* | Lib1-A73T | GGGAGCCTCCGGAGAAGCCCCGGGCCGA |
| *E06* | Lib1-C74G | GGCTGCCTCCGGAGAAGCCCCGGGCCGA |
| *F06* | Lib1-C75G | GCGTGCCTCCGGAGAAGCCCCGGGCCGA |
| *G06* | Lib1-C76G | CGGTGCCTCCGGAGAAGCCCCGGGCCGA |
| | **PLATE 3** | |
| **WellPosition** | **Name** | **Sequence** |
| *A01* | Lib1-WT | TCCGGAGGCACCCACTGCCACCGCGAAGAGTTGGGCTCTGTCAGCCGCGGGA |
| *C05* | Lib1-T64A | ACCGGAGGCACCCACTGCCACCGCGAAGAGTTGGGCTCTGTCAGCCGCGGGA |
| *D05* | Lib1-C65G | TGCGGAGGCACCCACTGCCACCGCGAAGAGTTGGGCTCTGTCAGCCGCGGGA |
| *E05* | Lib1-C66G | TCGGGAGGCACCCACTGCCACCGCGAAGAGTTGGGCTCTGTCAGCCGCGGGA |
| *F05* | Lib1-G67C | TCCCGAGGCACCCACTGCCACCGCGAAGAGTTGGGCTCTGTCAGCCGCGGGA |
| *G05* | Lib1-G68C | TCCGCAGGCACCCACTGCCACCGCGAAGAGTTGGGCTCTGTCAGCCGCGGGA |
| *H05* | Lib1-A69T | TCCGGTGGCACCCACTGCCACCGCGAAGAGTTGGGCTCTGTCAGCCGCGGGA |
| *A06* | Lib1-G70C | TCCGGACGCACCCACTGCCACCGCGAAGAGTTGGGCTCTGTCAGCCGCGGGA |
| *B06* | Lib1-G71C | TCCGGAGCCACCCACTGCCACCGCGAAGAGTTGGGCTCTGTCAGCCGCGGGA |
| *C06* | Lib1-C72G | TCCGGAGGGACCCACTGCCACCGCGAAGAGTTGGGCTCTGTCAGCCGCGGGA |
| *D06* | Lib1-A73T | TCCGGAGGCTCCCACTGCCACCGCGAAGAGTTGGGCTCTGTCAGCCGCGGGA |
| *E06* | Lib1-C74G | TCCGGAGGCAGCCACTGCCACCGCGAAGAGTTGGGCTCTGTCAGCCGCGGGA |
| *F06* | Lib1-C75G | TCCGGAGGCACGCACTGCCACCGCGAAGAGTTGGGCTCTGTCAGCCGCGGGA |
| *G06* | Lib1-C76G | TCCGGAGGCACCGACTGCCACCGCGAAGAGTTGGGCTCTGTCAGCCGCGGGA |
| *H06* | Lib1-A77T | TCCGGAGGCACCCTCTGCCACCGCGAAGAGTTGGGCTCTGTCAGCCGCGGGA |
| *A07* | Lib1-C78G | TCCGGAGGCACCCAGTGCCACCGCGAAGAGTTGGGCTCTGTCAGCCGCGGGA |
| *B07* | Lib1-T79A | TCCGGAGGCACCCACAGCCACCGCGAAGAGTTGGGCTCTGTCAGCCGCGGGA |
| *C07* | Lib1-G80C | TCCGGAGGCACCCACTCCCACCGCGAAGAGTTGGGCTCTGTCAGCCGCGGGA |
| *D07* | Lib1-C81G | TCCGGAGGCACCCACTGGCACCGCGAAGAGTTGGGCTCTGTCAGCCGCGGGA |
| *E07* | Lib1-C82G | TCCGGAGGCACCCACTGCGACCGCGAAGAGTTGGGCTCTGTCAGCCGCGGGA |
| *F07* | Lib1-A83T | TCCGGAGGCACCCACTGCCTCCGCGAAGAGTTGGGCTCTGTCAGCCGCGGGA |
| *G07* | Lib1-C84G | TCCGGAGGCACCCACTGCCAGCGCGAAGAGTTGGGCTCTGTCAGCCGCGGGA |
| *H07* | Lib1-C85G | TCCGGAGGCACCCACTGCCACGGCGAAGAGTTGGGCTCTGTCAGCCGCGGGA |
| *A08* | Lib1-G86C | TCCGGAGGCACCCACTGCCACCCCGAAGAGTTGGGCTCTGTCAGCCGCGGGA |
| *B08* | Lib1-C87G | TCCGGAGGCACCCACTGCCACCGGGAAGAGTTGGGCTCTGTCAGCCGCGGGA |
| *C08* | Lib1-G88C | TCCGGAGGCACCCACTGCCACCGCCAAGAGTTGGGCTCTGTCAGCCGCGGGA |

| | | |
|---|---|---|
| *D08* | Lib1-A89T | TCCGGAGGCACCCACTGCCACCGCGTAGAGTTGGGCTCTGTCAGCCGCGGGA |
| *E08* | Lib1-A90T | TCCGGAGGCACCCACTGCCACCGCGATGAGTTGGGCTCTGTCAGCCGCGGGA |
| *F08* | Lib1-G91C | TCCGGAGGCACCCACTGCCACCGCGAACAGTTGGGCTCTGTCAGCCGCGGGA |
| *G08* | Lib1-A92T | TCCGGAGGCACCCACTGCCACCGCGAAGTGTTGGGCTCTGTCAGCCGCGGGA |
| *H08* | Lib1-G93C | TCCGGAGGCACCCACTGCCACCGCGAAGACTTGGGCTCTGTCAGCCGCGGGA |
| *A09* | Lib1-T94A | TCCGGAGGCACCCACTGCCACCGCGAAGAGATGGGCTCTGTCAGCCGCGGGA |
| *B09* | Lib1-T95A | TCCGGAGGCACCCACTGCCACCGCGAAGAGTAGGGCTCTGTCAGCCGCGGGA |
| *C09* | Lib1-G96C | TCCGGAGGCACCCACTGCCACCGCGAAGAGTTCGGCTCTGTCAGCCGCGGGA |
| *D09* | Lib1-G97C | TCCGGAGGCACCCACTGCCACCGCGAAGAGTTGCGCTCTGTCAGCCGCGGGA |
| *E09* | Lib1-G98C | TCCGGAGGCACCCACTGCCACCGCGAAGAGTTGGCCTCTGTCAGCCGCGGGA |
| *F09* | Lib1-C99G | TCCGGAGGCACCCACTGCCACCGCGAAGAGTTGGGGTCTGTCAGCCGCGGGA |
| *G09* | Lib1-T100A | TCCGGAGGCACCCACTGCCACCGCGAAGAGTTGGGCACTGTCAGCCGCGGGA |
| *H09* | Lib1-C101G | TCCGGAGGCACCCACTGCCACCGCGAAGAGTTGGGCTGTGTCAGCCGCGGGA |
| *A10* | Lib1-T102A | TCCGGAGGCACCCACTGCCACCGCGAAGAGTTGGGCTCAGTCAGCCGCGGGA |
| *B10* | Lib1-G103C | TCCGGAGGCACCCACTGCCACCGCGAAGAGTTGGGCTCTCTCAGCCGCGGGA |
| *C10* | Lib1-T104A | TCCGGAGGCACCCACTGCCACCGCGAAGAGTTGGGCTCTGACAGCCGCGGGA |
| *D10* | Lib1-C105G | TCCGGAGGCACCCACTGCCACCGCGAAGAGTTGGGCTCTGTGAGCCGCGGGA |
| *E10* | Lib1-A106T | TCCGGAGGCACCCACTGCCACCGCGAAGAGTTGGGCTCTGTCTGCCGCGGGA |
| *F10* | Lib1-G107C | TCCGGAGGCACCCACTGCCACCGCGAAGAGTTGGGCTCTGTCACCCGCGGGA |
| *G10* | Lib1-C108G | TCCGGAGGCACCCACTGCCACCGCGAAGAGTTGGGCTCTGTCAGGCGCGGGA |
| *H10* | Lib1-C109G | TCCGGAGGCACCCACTGCCACCGCGAAGAGTTGGGCTCTGTCAGCGGCGGGA |
| *A11* | Lib1-G110C | TCCGGAGGCACCCACTGCCACCGCGAAGAGTTGGGCTCTGTCAGCCCCGGGA |
| *B11* | Lib1-C111G | TCCGGAGGCACCCACTGCCACCGCGAAGAGTTGGGCTCTGTCAGCCGGGGGA |
| *C11* | Lib1-G112C | TCCGGAGGCACCCACTGCCACCGCGAAGAGTTGGGCTCTGTCAGCCGCCGGA |
| *D11* | Lib1-G113C | TCCGGAGGCACCCACTGCCACCGCGAAGAGTTGGGCTCTGTCAGCCGCGCGA |
| *E11* | Lib1-G114C | TCCGGAGGCACCCACTGCCACCGCGAAGAGTTGGGCTCTGTCAGCCGCGGCA |
| | **PLATE 4** | |
| **WellPosition** | **Name** | **Sequence** |
| *A01* | Lib1-WT | GTTGTTGTTGTTGTTTCTTTTGTTGTAAGTACTCCTTACTTTGTTTTCCCGCGGCTGA |
| *C10* | Lib1-T104A | GTTGTTGTTGTTGTTTCTTTTGTTGTAAGTACTCCTTACTTTGTTTTCCCGCGGCTGT |
| *D10* | Lib1-C105G | GTTGTTGTTGTTGTTTCTTTTGTTGTAAGTACTCCTTACTTTGTTTTCCCGCGGCTCA |
| *E10* | Lib1-A106T | GTTGTTGTTGTTGTTTCTTTTGTTGTAAGTACTCCTTACTTTGTTTTCCCGCGGCAGA |
| *F10* | Lib1-G107C | GTTGTTGTTGTTGTTTCTTTTGTTGTAAGTACTCCTTACTTTGTTTTCCCGCGGGTGA |
| *G10* | Lib1-C108G | GTTGTTGTTGTTGTTTCTTTTGTTGTAAGTACTCCTTACTTTGTTTTCCCGCGCCTGA |
| *H10* | Lib1-C109G | GTTGTTGTTGTTGTTTCTTTTGTTGTAAGTACTCCTTACTTTGTTTTCCCGCCGCTGA |
| *A11* | Lib1-G110C | GTTGTTGTTGTTGTTTCTTTTGTTGTAAGTACTCCTTACTTTGTTTTCCCGGGGCTGA |
| *B11* | Lib1-C111G | GTTGTTGTTGTTGTTTCTTTTGTTGTAAGTACTCCTTACTTTGTTTTCCCCCGGCTGA |
| *C11* | Lib1-G112C | GTTGTTGTTGTTGTTTCTTTTGTTGTAAGTACTCCTTACTTTGTTTTCCGGCGGCTGA |
| *D11* | Lib1-G113C | GTTGTTGTTGTTGTTTCTTTTGTTGTAAGTACTCCTTACTTTGTTTTCGCGCGGCTGA |
| *E11* | Lib1-G114C | GTTGTTGTTGTTGTTTCTTTTGTTGTAAGTACTCCTTACTTTGTTTTGCCGCGGCTGA |
